# Fatty acid-binding proteins and fatty acid synthase influence glial reactivity and promote the formation of Müller glia-derived progenitor cells in the avian retina

**DOI:** 10.1101/2021.08.19.456977

**Authors:** Warren A. Campbell, Allen Tangeman, Heithem M. El-Hodiri, Evan C. Hawthorn, Maddie Hathoot, Thanh Hoang, Seth Blackshaw, Andy J. Fischer

## Abstract

The capacity for retinal regeneration varies greatly across vertebrates species. A recent comparative epigenetic and transcriptomic investigation of Müller glial (MG) in the retinas of fish, birds and mice revealed that Fatty Acid Binding Proteins (FABPs) are among the most highly up-regulated genes in activated chick MG (Hoang et al., 2020). Herein we provide an in-depth follow-up investigation to describe patterns of expression and how FABPs and fatty acid synthase (FASN) influence glial cells in the chick retina. During development, *FABP7* is highly expressed by embryonic retinal progenitor cells (eRPCs) and maturing MG, whereas *FABP5* is gradually up-regulated in maturing MG and remains elevated in mature glial cells. *PMP2* (FABP8) is expressed by oligodendrocytes and *FABP5* is expressed by non-astrocytic inner retinal glial cells, and both of these FABPs are significantly up-regulated in activated MG in damaged or growth factor-treated retinas. In addition to suppressing the formation of MGPCs, we find that FABP-inhibition suppressed the accumulation of proliferating microglia, although the microglia appeared highly reactive. scRNA-seq analyses of cells treated with FABP-inhibitor revealed distinct changes in patterns of expression suggesting that FABPs are involved in the transitions of MG from a resting state to a reactive state and conversion from reactive MG to MGPCs. Inhibition of FABPs in undamaged retinas had a significant impact upon the transcriptomic profiles of MG, with up-regulation of genes associated with gliogenesis, decreases in genes associated with neurogenesis, and suppression of the ability of MG to become MGPCs. scRNA-seq analyses of microglia indicated that FABP inhibition enhances gene modules related to reactivity, proliferation and cytokine signaling. We find that the proliferation of retinal progenitors in the circumferential marginal zone (CMZ) is unaffected by FABP-inhibitor. Upstream of FABP activity, we inhibited FASN in damaged retinas, which reduced numbers of dying cells, increased the proliferation of microglia, and potently suppressed the formation MGPCs in damaged retinas. We conclude that the activity of FASN and FABPs are required early during the formation of proliferating MGPCs. Fatty acid metabolism and cell signaling involving fatty acids are important in regulating glial homeostasis in the retina, and the dedifferentiation and proliferation of microglia and MGPCs.

## Introduction

The process of retinal regeneration varies greatly across vertebrate species. In the fish, retinal neuronal regeneration is a robust process that restores functional cells and visual acuity following injury, whereas this process is far less robust in birds and absent in mammals (Hitchcock and Raymond, 1992; Karl et al., 2008; Raymond, 1991). Müller glia (MG) have been identified as the cell of origin for progenitors in regenerating retinas (Bernardos et al., 2007; Fausett and Goldman, 2006; Fausett et al., 2008; Fischer and Reh, 2001; Ooto et al., 2004). In normal healthy retinas, MG are the predominant type of support cell that provide structural, metabolic, visual cycle, and synaptic support (Reichenbach and Bringmann, 2013). In response to damage, certain growth factors or drug treatments, MG can be stimulated to become reactive, de-differentiate, up-regulate progenitor-related genes, re-enter the cell cycle and produce progeny that differentiate as neurons (Fischer and Bongini, 2010; Gallina et al., 2014a; Wan and Goldman, 2016).

In mammalian retinas, significant stimulation such as forced expression of *Ascl1*, inhibition of histone deacetylases and neuronal damage is required to reprogram MG into progenitor-like cells that produce a few new neurons (Jorstad et al., 2017; Pollak et al., 2013; Ueki et al., 2015). Alternatively, deletion of *Nfia, Nfib* and *Nfix* in mature MG combined with retinal damage and treatment with insulin+FGF2 results in reprogramming of MG into cells that resemble inner retinal neurons (Hoang et al., 2020). Blockade of Hippo-signaling via forced expression of degradation-resistant YAP1 drives the proliferation of mature MG in the mouse retina (Hamon et al., 2019; Rueda et al., 2019), but it remains unknown whether any of the progeny differentiate as neurons. In addition, viral delivery of reporters, β-catenin, *Otx2*, *Crx* and *Nrl* may reprogram MG into photoreceptors (Yao et al., 2018), but there are concerns that the viral vectors and mini-promoters used in these studies are prone to leaky expression in neurons (Blackshaw and Sanes, 2021). In the chick retina, MG readily reprogram into progenitor-like cells that proliferate, but the progeny have a limited capacity to differentiate as neurons (Fischer and Reh, 2001; Fischer and Reh, 2003). Understanding the mechanisms that regulate the formation of MGPCs and neuronal differentiation of progeny is important to harnessing the regenerative potential of MG in warm-blooded vertebrates.

Fatty acid synthesis, metabolism and signaling are likely to be key components of regulating neuronal progenitor cells. Fatty Acid Binding Proteins (FABPs) are cytosolic lipid-binding proteins, that mediate fatty acid metabolism and cell-signaling, and have highly conserved primary and tertiary structures across species from *Drosophila* to humans (Hanhoff et al., 2002; Smathers and Petersen, 2011). FABPs are known to bind to poly-unsaturated fatty acids (PUFAs) including arachidonic acid and docosahexaenoic acid and have been shown to regulate signal transduction, neurotransmission, proliferation, differentiation, and cell migration (Allen et al., 2007; Dawson and Xia, 2012; Tripathi et al., 2017; Yamashima, 2012). Very little is known about the cellular mechanisms and patterns of expression of FABPs in the retina. In mammals, FABP3, 5, and 7 have been identified in the brain, retina, and radial glia with demonstrated roles in differentiation and cell fate determination (Owada, 2008; Sellner, 1993; Sellner et al., 1995) llen et al., 2007; Dawson and Xia, 2012; Tripathi et al., 2017; Yamashima, 2012). Further evidence indicates that FABPs in the CNS modulate endocannabinoid, Peroxisome Proliferator-Activated Receptor (PPAR), NF-kB, and CREB signaling (Bogdan et al., 2018; Peng et al., 2017; Tripathi et al., 2017; Yamashima, 2012). NF-kB has been implicated as a key signaling “hub” that suppresses the formation of MGPCs in chicks and mice, but not zebrafish (Hoang et al., 2020; Palazzo et al., 2020).

We have previously reported that *FABP5* and *PMP2* are highly up-regulated in MG in NMDA-damaged retinas, and that inhibition of FABPs potently suppresses the formation of proliferating MGPCs (Hoang et al., 2020). However, few details are known about the mechanisms by which FABPs act to influence the formation of MGPCs, coordinate with other cell-signaling pathways, influence the reactivity of microglia, and induce changes in gene expression following FABP-inhibition. Accordingly, this study investigates how FABPs and Fatty Acid Synthase (FASN) influence reprogramming of MG into MGPCs and analyze transcriptomic changes downstream of FABP inhibition in damaged and growth factor-treated retinas in the chick model system.

## Methods and Materials

### Animals

The animals approved for use in these experiments was in accordance with the guidelines established by the National Institutes of Health and IACUC at The Ohio State University. Newly hatched P0 wildtype leghorn chicks (*Gallus gallus domesticus*) were obtained from Meyer Hatchery (Polk, Ohio). Post-hatch chicks were maintained in a regular diurnal cycle of 12 hours light, 12 hours dark (8:00 AM-8:00 PM). Chicks were housed in stainless-steel brooders at 25°C and received water and Purina^tm^ chick starter *ad libitum*.

Fertilized eggs were obtained from the Michigan State University, Department of Animal Science. Eggs were incubated at a constant 37.5°C, with a 1hr period at room temperature with a cool-down period every 24hrs. The eggs were rocked every 45 minutes and held at a constant relative humidity of 45%. Embryos were harvested at various time points after incubation and staged according to guidelines established by Hamburger and Hamilton (Hamburger and Hamilton, 1992).

### Intraocular injections

Chicks were anesthetized with 2.5% isoflurane mixed with oxygen from a non-rebreathing vaporizer. The technical procedures for intraocular injections were performed as previously described (Fischer et al., 1998). With all injection paradigms, both pharmacological and vehicle treatments were administered to the right and left eye respectively. Compounds were injected in 20 μl sterile saline with 0.05 mg/ml bovine serum albumin added as a carrier. Compounds included: NMDA (38.5nmol or 154 µg/dose; Millipore Sigma), FGF2 (250 ng/dose; R&D systems), BMS309403 (Millipore Sigma), C75 (Millipore Sigma), G28UCM (Millipore Sigma). 5-Ethynyl-2ʼ-deoxyuridine (EdU, ThermoFischer) was injected into the vitreous chamber to label proliferating cells. Injection paradigms are included in each figure.

### Single Cell RNA sequencing of retinas

Retinas were obtained from embryonic and hatched chicks. Retinas were dissociated in a 0.25% papain solution in Hank’s balanced salt solution (HBSS), pH = 7.4, for 30 minutes, and suspensions were frequently triturated. The dissociated cells were passed through a sterile 70µm filter to remove large particulate debris. Dissociated cells were assessed for viability (Countess II; Invitrogen) and cell-density diluted to 700 cell/µl. Each single cell cDNA library was prepared for a target of 10,000 cells per sample. The cell suspension and Chromium Single Cell 3’ V2 or V3 reagents (10X Genomics) were loaded onto chips to capture individual cells with individual gel beads in emulsion (GEMs) using the 10X Chromium Cell Controller. cDNA and library amplification and for an optimal signal was 12 and 10 cycles respectively. Sequencing was conducted on Illumina HiSeq2500 (Genomics Resource Core Facility, John’s Hopkins University), or Novaseq6000 (Novogene) using 150 paired-end reads. Fasta sequence files were de-multiplexed, aligned, and annotated using the chick ENSMBL database (GRCg6a, Ensembl release 94) by using 10X Cell Ranger software. Gene expression was counted using unique molecular identifier bar codes and gene-cell matrices were constructed. Using Seurat toolkits Uniform Manifold Approximation and Projection for Dimension Reduction (UMAP) plots were generated from aggregates of multiple scRNA-seq libraries (Butler et al., 2018; Satija et al., 2015). Seurat was used to construct gene lists for differentially expressed genes (DEGs), violin/scatter plots and dot plots. Significance of difference in violin/scatter plots was determined using a Wilcoxon Rank Sum test with Bonferroni correction. Monocle was used to construct unguided pseudotime trajectories and scatter plotters for MG and MGPCs across pseudotime (Qiu et al., 2017a; Qiu et al., 2017b; Trapnell et al., 2012).

SingleCellSignalR was used to assess potential ligand-receptor interactions between cells within scRNA-seq datasets (Cabello-Aguilar et al., 2020). Genes that were used to identify different types of retinal cells included the following: (1) Müller glia: *GLUL, VIM, SCL1A3, RLBP1*, (2) MGPCs: *PCNA, CDK1, TOP2A, ASCL1*, (3) microglia: *C1QA, C1QB, CCL4, CSF1R, TMEM22*, (4) ganglion cells: *THY1, POU4F2, RBPMS2, NEFL, NEFM*, (5) amacrine cells: *GAD67, CALB2, TFAP2A*, (6) horizontal cells: *PROX1, CALB2, NTRK1*, (7) bipolar cells: *VSX1, OTX2, GRIK1, GABRA1*, and (7) cone photoreceptors: *CALB1, GNAT2, OPN1LW*, and (8) rod photoreceptors: *RHO, NR2E3, ARR1.* Gene Ontology (GO) enrichment analysis was performed using ShinyGO V0.65 (http://bioinformatics.sdstate.edu/go/). scRNA-seq libraries can be queried at https://proteinpaint.stjude.org/F/2019.retina.scRNA.html or gene-cell matricies downloaded from GitHub at https://github.com/fischerlab3140/scRNAseq_libraries

### Fixation, sectioning and immunocytochemistry

Retinal tissue samples were formaldehyde fixed, sectioned, and labeled via immunohistochemistry as described previously (Fischer et al., 2006; Ghai et al., 2009). Antibody dilutions and commercial sources for images used in this study are described in table 1. Observed labeling was not due to off-target labeling of secondary antibodies or tissue auto-fluorescence because sections incubated exclusively with secondary antibodies were devoid of fluorescence. Secondary antibodies utilized include donkey-anti-goat-Alexa488/568, goat-anti-rabbit-Alexa488/568/647, goat-anti-mouse-Alexa488/568/647, goat-anti-rat-Alexa488 (Life Technologies) diluted to 1:1000 in PBS and 0.2% Triton X-100.

**Table 1.**
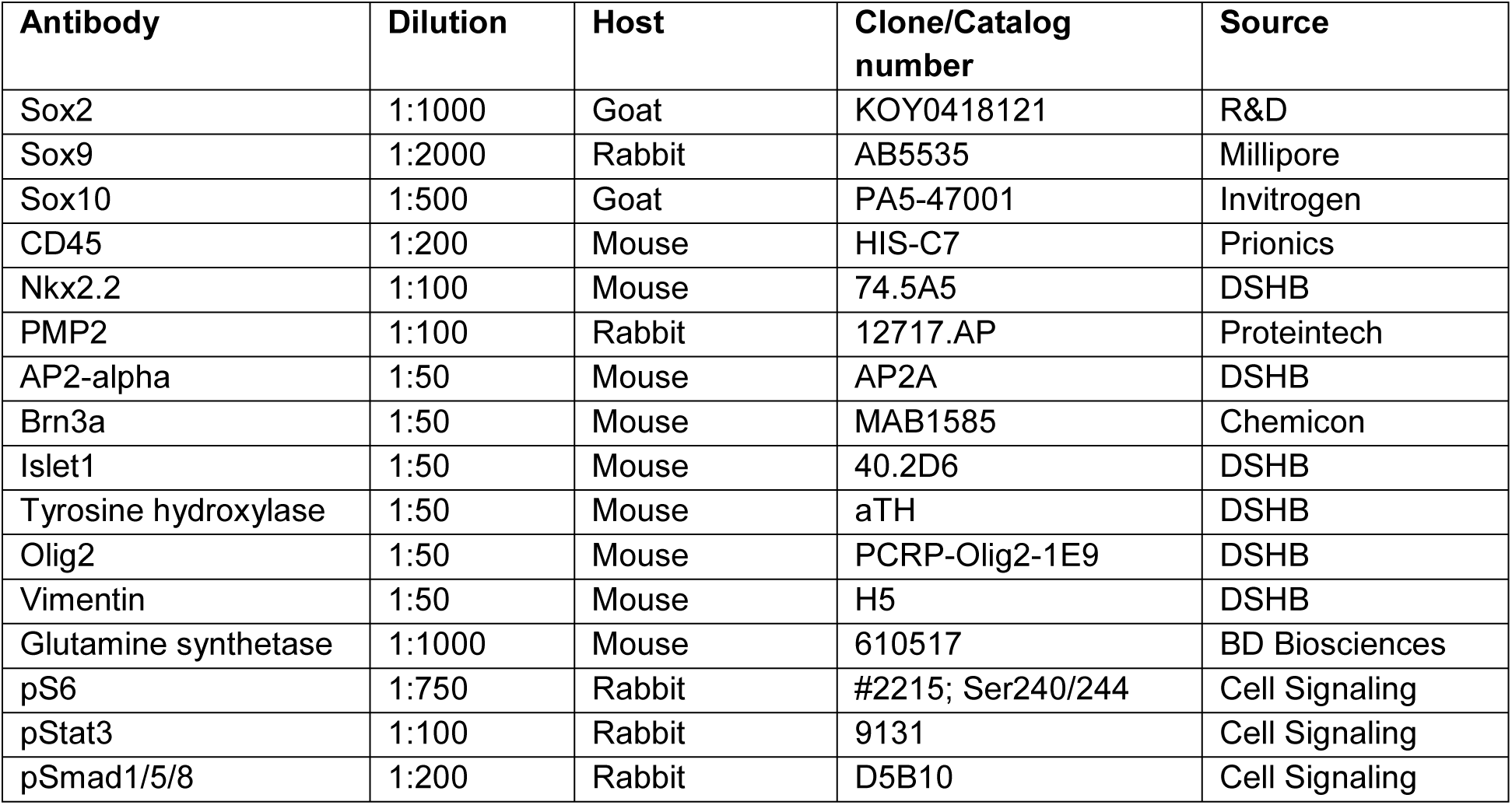
List of antibodies, working dilution, clone/catalog number and source.

| Antibody | Dilution | Host | Clone/Catalog number | Source |
| --- | --- | --- | --- | --- |
| Sox2 | 1:1000 | Goat | KOY0418121 | R&D |
| Sox9 | 1:2000 | Rabbit | AB5535 | Millipore |
| Sox10 | 1:500 | Goat | PA5-47001 | Invitrogen |
| CD45 | 1:200 | Mouse | HIS-C7 | Prionics |
| Nkx2.2 | 1:100 | Mouse | 74.5A5 | DSHB |
| PMP2 | 1:100 | Rabbit | 12717.AP | Proteintech |
| AP2-alpha | 1:50 | Mouse | AP2A | DSHB |
| Brn3a | 1:50 | Mouse | MAB1585 | Chemicon |
| Islet1 | 1:50 | Mouse | 40.2D6 | DSHB |
| Tyrosine hydroxylase | 1:50 | Mouse | aTH | DSHB |
| Olig2 | 1:50 | Mouse | PCRP-Olig2-1E9 | DSHB |
| Vimentin | 1:50 | Mouse | H5 | DSHB |
| Glutamine synthetase | 1:1000 | Mouse | 610517 | BD Biosciences |
| pS6 | 1:750 | Rabbit | #2215; Ser240/244 | Cell Signaling |
| pStat3 | 1:100 | Rabbit | 9131 | Cell Signaling |
| pSmad1/5/8 | 1:200 | Rabbit | D5B10 | Cell Signaling |

### Labeling for EdU

For the detection of nuclei that incorporated EdU, immunolabeled sections were fixed in 4% formaldehyde in 0.1M PBS pH 7.4 for 5 minutes at room temperature. Samples were washed for 5 minutes with PBS, permeabilized with 0.5% Triton X-100 in PBS for 1 minute at room temperature and washed twice for 5 minutes in PBS. Sections were incubated for 30 minutes at room temperature in a buffer consisting of 100 mM Tris, 8 mM CuSO_4_, and 100 mM ascorbic acid in dH_2_O. The Alexa Fluor 568 Azide (Thermo Fisher Scientific) was added to the buffer at a 1:100 dilution.

### Terminal deoxynucleotidyl transferase dUTP nick end labeling (TUNEL)

The TUNEL assay was implemented to identify dying cells by imaging fluorescent labeling of double stranded DNA breaks in nuclei. The *In Situ* Cell Death Kit (TMR red; Roche Applied Science) was applied to fixed retinal sections as per the manufacturer’s instructions.

### Photography, measurements, cell counts and statistics

Microscopy images of retinal sections were captured with the Leica DM5000B microscope with epifluorescence and the Leica DC500 digital camera. High resolution confocal images were obtained with a Leica SP8 available in The Department of Neuroscience Imaging Facility at The Ohio State University. Representative images are modified to have enhanced color, brightness, and contrast for improved clarity using Adobe Photoshop. In EdU proliferation assays, a fixed region of retina was counted and average numbers of Sox2 and EdU co-labeled cells. The retinal region selected for investigation was standardized between treatment and control groups to reduce variability and improve reproducibility.

Similar to previous reports (Ghai et al., 2009), immunofluorescence intensity was quantified by using Image J (NIH). Identical illumination, microscope, and camera settings were used to obtain images for quantification. Retinal areas were sampled from digital images. These areas were randomly sampled over the inner nuclear layer (INL). MS Excel and GraphPad Prism 6 were used for statistical analyses. Measurement for immunofluorescence in the nuclei of MG/MGPCs were made by from single optical confocal sections by selecting the total area of pixel values above threshold (≥70) for Sox2 or Sox9 immunofluorescence (in the red channel) and copying nuclear labeling from only MG (in the green channel). Measurements of pS6 immunofluorescence were made for a fixed, cropped area (14,000 µm^2^) of INL, OPL and ONL. Measurements were made for regions containing pixels with intensity values of 70 or greater (0 = black and 255 = saturated). The total area was calculated for regions with pixel intensities above threshold. The intensity sum was calculated as the total of pixel values for all pixels within threshold regions.

For statistical evaluation of differences in treatments, a two-tailed paired *t*-test was applied for intra-individual variability where each biological sample also served as its own control. For two treatment groups comparing inter-individual variability, a two-tailed unpaired *t*-test was applied. For multivariate analysis, an ANOVA with the associated Tukey Test was used to evaluate any significant differences between multiple groups.

## Results

### Expression of *FABPs* during embryonic retinal development

scRNA-seq libraries were established for retinal cells at embryonic day 5 (E5), E8, E12, and E15. These libraries yielded 22,698 cells after filtering to exclude doublets, cells with low UMI/genes per cell, and high mitochondrial gene-content. UMAP plots of aggregate libraries formed clusters of retinal cells that correlated to developmental stage and cell type (Fig. 1a). Types of cells were identified based on expression of well-established markers. Retinal progenitor cells (RPCs) from E5 and E8 retinas were identified by expression of *ASCL1*, C*DK1*, and *TOP2A*. (Fig. 1b). Maturing MG were identified by expression of *GLUL*, *RLBP1* and *SLC1A3* (Fig. 1b). *FABP5* and *FABP7* were expressed by different types of developing retinal cells at different stages of development, whereas *PMP2* (FABP8) was not widely expressed by embryonic retinal cells (Fig. 1c). *FABP5* was predominantly expressed by maturing bipolar and amacrine cells (Figs. 1c,d). By comparison, *FAPB7* was predominantly expressed by early retinal progenitor cells (eRPCs) from E5 and E8 retinas and at elevated levels in immature MG at E8, whereas levels decreased in maturing MG at E12 and E15 (Figs.1c,d,f). *FABP7* was also expressed by maturing bipolar and amacrine cells from E8 retinas (Figs. 1c,d).

**Figure 1.**
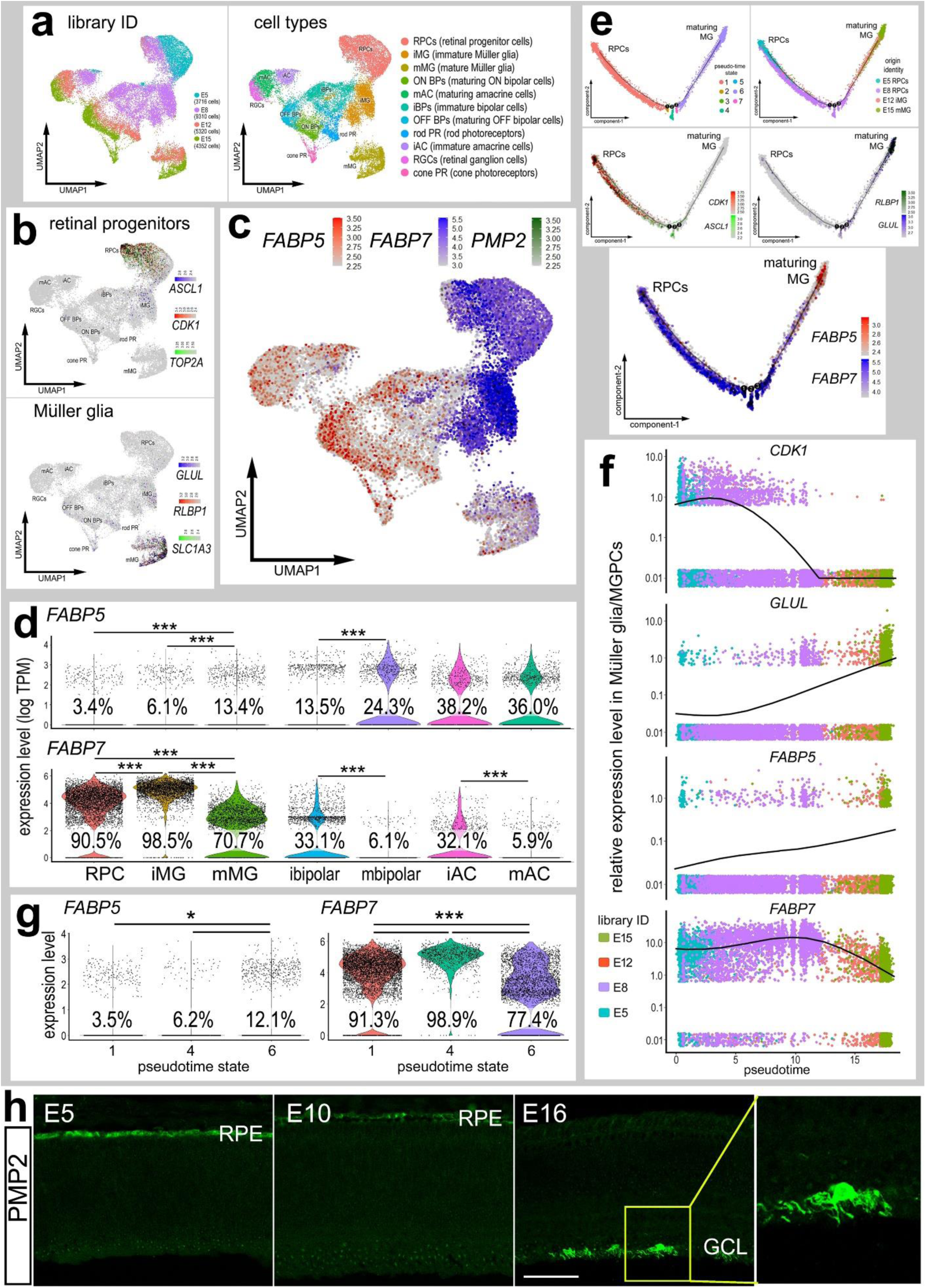
Expression of *FABP5*, *FABP7* and *PMP2* in embryonic chick retina. scRNA-seq libraries were generated from embryonic retinal cells at four stages of development (E5, E8, E12, E15) (**a**). UMAP clustered were generated to identify cell types and probe for gene expression (**b**). Cells were identified based on expression of cell-distinguishing genes (**a**,**b**). FABP isoforms were plotted in a heatmap, with cells expressing 2+ genes denoted in black (**c**).The expression of FABPs in different maturing cell types is represented in violin plots (**d**). MG were also plotted in pseudotime denoting their transition from an immature progenitor cell to mature glial cell (**e**). *FABP5* and *FABP7* expression fluctuated during this transition as denoted by pseudotime scatter plots (**f**) and pseudotime state violin plots (**g**). Significant difference (*p<0.01, **p<0.0001, ***p<<0.0001) was determined by using a Wilcox rank sum with Bonferoni correction. RPC – retinal progenitor cell, MG – Müller glia, iMG – immature Müller glia, mMG - mature Müller glia.

To further assess patterns of expression we re-embedding of eRPCs and MG to order cells across pseudotime in an unsupervised manner. Pseudotime-ordering revealed a continuum of cells with early eRPCs and maturing MG at opposite ends of the trajectory (Fig. 1e). Across pseudotime levels of *GLUL* increased, while levels of *CDK1* decreased (Fig. 1e,f). Like *GLUL*, the expression of *FABP5* increases from retinal progenitors to maturing MG (Fig. 1e-g). By contrast, levels of *FABP7* were relatively high in eRPCs, increased in differentiating immature MG, and decreased in MG during later stages of development (Figs. 1e-g). Immunolabeling for PMP2 confirmed findings from scRNA-seq. PMP2 was not expressed in the developing retina until E16, when PMP2-immunofluorescence was detected near the vitread surface of the retina in cells that resembled oligodendrocytes (Fig. 1h). These findings indicate that *FABP5* and *FABP7* are dynamically and highly expressed in eRPCs and MG through the course of embryonic development.

### Expression of FABPs in damage to the chick retina

We sought to provide a detailed description of patterns of expression of FABPs in the retinas of normal and hatched chick by using scRNA-seq. Libraries were aggregated for retinal cells obtained from control and NMDA-damaged retinas at different time points (24, 48 and 72 hrs) after treatment for a total of 57,230 cells (Fig. 2a). We have previously used these chick scRNA-seq databases to compare MG and MGPCs across fish, chick and mouse model systems (Hoang et al., 2020) and characterize expression patterns of genes related to NFkB-signaling (Palazzo et al., 2020), midkine-signaling (Campbell et al., 2021), matrix metalloproteases (Campbell et al., 2019), and endocannabinoid-signaling (Campbell et al., 2021b). UMAP-clustered cells were identified based on well-established patterns of expression (Fig. 2a,b). Resting MG formed a discrete cluster of cells and expressed high levels of *GLUL, RLBP1* and *SLC1A3* (Fig. 2a,b). After damage, MG down-regulate markers of mature glia as they transition into reactive glial cells and into progenitor-like cells that up-regulate *TOP2A, CDK1* and *ESPL1* (Fig. 2a,b). *FABP5* and *PMP2* were expressed at low levels in relatively few resting MG in undamaged retina, whereas *FABP7* was widely expressed by the majority of resting MG (Fig. 2c,d). *FABP7* and *PMP2* were detected in oligodendrocytes and Non-astrocytic Inner Retinal Glia (NIRGs). NIRG cells are a distinct type of glial cell that has been described in the retinas of birds (Rompani and Cepko 2010; Fischer et al., 2010) and some types of reptiles (Todd et al., 2016b). Following NMDA-induced retinal damage, levels of *FABP5, FABP7* and *PMP2* were significantly increased in activated MG at 24 hrs (Fig. 2c,d). Levels of *FABP5 a*nd *PMP2* were significantly reduced in activated MG at 48hrs and 72hrs, but remained elevated in proliferating MGPCs (Fig. 2c,d).

**Figure 2.**
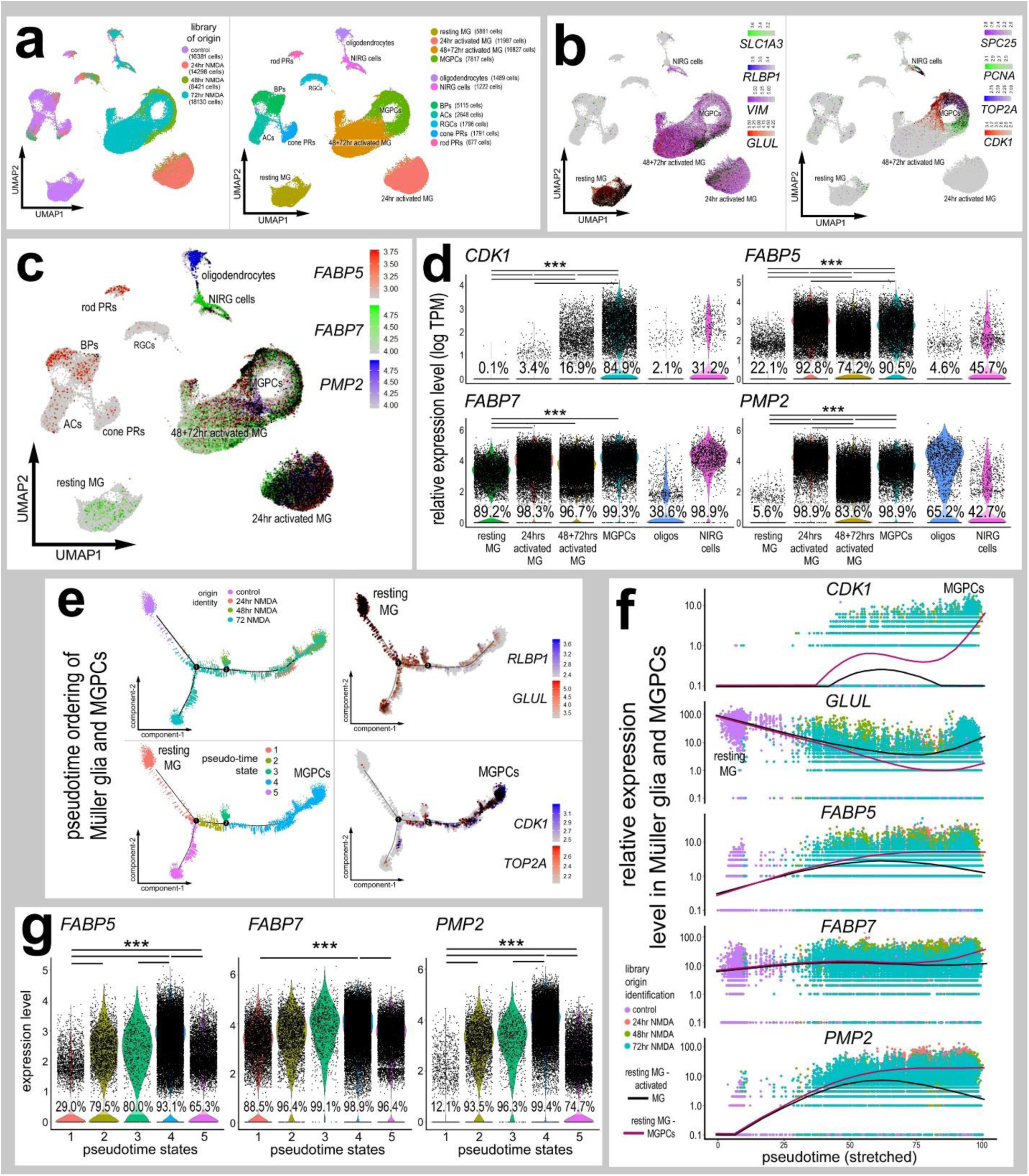
Chick MG of damaged retinas express FABP isoforms preferentially during the MGPC transition. scRNA-seq libraries of time points after NMDA damage were aggregated and clustered with UMAP to identify unique MG clusters transitioning into MGPCs (**a,b**). The levels of FABP7, FABP5, PMP2 in MG clusters, oligodendrocytes, and nonastrocytic inner retinal glia (NIRG) cells are represented by violin plots (**d**). The response of MG to damage was modelled in pseudotime, indicating a divergent response between glial reactivity and de-differentiated MGPC (**e**). The expression levels of FABPs are compared between a “reactive” and “reprogramming” response by MG. FABP expression are shown in violin plots of MG in different transitional states (**f**). These divergent transitions are shown on the pseudotime scatter plot with the reactive and reprogramming branch are denoted by the black and red lines respectively (**g**). Each dot in violin and scatter plots represents an individual cell. Significant changes (*p<0.1, **p<0.0001, ***p<<0.0001) in levels were determined by using a Wilcox rank sum with Bonferroni correction.

We re-ordered MG along pseudotime trajectories to better assess changes in expression of *FABPs* during the transition of MG to progenitor cells. We identified 5 pseudotime states and a trajectory that included distinct branches for resting Müller glia (state 1), activated Müller glia (state 5), transitional Müller glia (states 2,3), and MGPCs (state 4) (Fig. 2e). Dimensional reduction to a single pseudotime axis placed resting Müller glia (high levels of *GLUL*) to the left and highly activated MG and MGPCs (high levels of *CDK1*) the far right of the pseudotime axis (Fig. 2e-f). The expression of *FABP5* and *PMP2* across pseudotime positively correlated with a transition toward an MGPC-phenotype, and inversely correlated to resting glial phenotypes (Fig. 2f,g). Levels of *FABP7* were not significantly different across most pseudotime states, with the exception of elevated levels in state 4 which is occupied by MGPCs, compared to levels in resting and activated MG (Fig. 2f,g).

We next assessed patterns of expression of FABPs in scRNA-seq libraries from time-points soon after NMDA at 3 and 12 hrs after treatment. These libraries were generated with newer, more sensitive reagents and did not integrate well with older libraries generated with less sensitive reagents. UMAP ordering of MG revealed distinct clusters of cells which closely match treatment groups and included resting MG, early activated MG, 3 groups of activated MG and 2 groups of MGPCs (supplemental Fig. 1a,b). Levels of *FABP5* were reduced, but expressed by a larger percentage of MG at 3hrs after NMDA, levels increased in activated MG, and further increased in MGPCs (supplemental Fig. 1e,h). *FABP7* was down-regulated in MG at 12hrs, but increased in MGPCs at 48hrs after NMDA (supplemental Fig. 1f,h). Levels of *PMP2* were not increased in MG until 12hrs after NMDA-treatment, and remained elevated in MGPCs through 48hrs after treatment (supplemental Fig. 1g,h).

### Immunolabeling for PMP2

We next sought to characterize PMP2 immunolabeling in normal retinas. We found that PMP2 immunofluorescence was observed in NIRG cells, but only in peripheral regions of retina (Fig. 3a). Although the NIRG cells were often observed in the proximal INL and had morphologies reminiscent of amacrine cells; these cells were negative for amacrine cell markers including AP2α, Islet1 and tyrosine hydroxylase (Fig. 3a). PMP2-immunolabeling was prevalent in oligodendrocytes extended processes into the NFL, and were negative for Glutamine Synthase (GS; Fig. 3b). The PMP2-positive oligodendrocytes where immunoreactive for Olig2 (Fig. 3c), Sox10 (Fig. 3g) and HuC/D (Fig. 3d). HuC/D is thought to be a neuron-specific marker, but appears to be up-regulated in PMP2+ oligodendrocytes after NMDA-treatment (Fig. 3d) and a few of these cells proliferate (accumulate EdU) in central and peripheral regions of the damaged retinas (Fig. 3e,f). NIRG cells were not immunoreactive for HuC/D at any time after NMDA-treatment. Further, we failed to observe PMP2 in NIRG cells (Sox2+/Nkx2.2+ cells in the IPL) in central regions of control or NMDA-damaged retinas (Fig. 3h,i). Collectively, these patterns of immunolabeling are consistent with scRNA-seq data for oligodendrocytes and NIRG cells and patterns of expression for *PMP2, SOX10, OILG2* and *ELAVL4* (HuD; Fig. 3j). Further, NIRG cells expressed *FABP5* and *FABP7*, whereas oligodendrocytes expressed *PMP2* (Fig. 3j).

**Figure 3.**
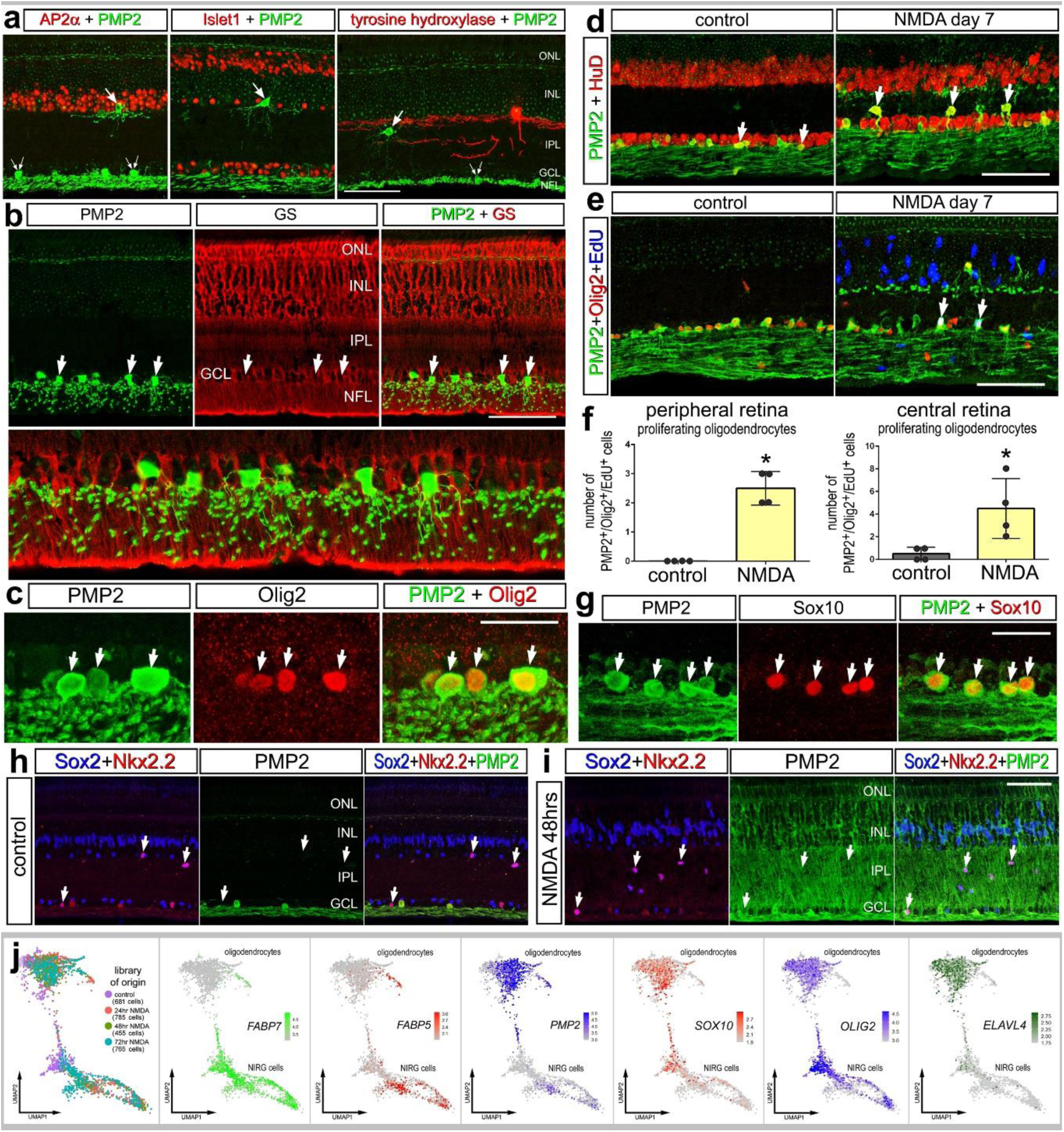
PMP2-immunoreactivity in chick retinas. Vertical sections of the retina were obtained from untreated eyes (**a**-**e**, **g**, **h**) and eyes injected with NMDA (**i**). Tissues were labeled with antibodies to PMP2 (green) and Ap2α (red; **a**), Islet1 (red; **a**), tyrosine hydroxylase (red; **a**), GS (red; **b**), Olig2 (red, **c,e**), HuC/D (red; **d**), EdU (blue; **g**), Sox10 (red; **g**), Sox2 (blue; **h,i**), and Nkx2.2 (red; **h,i**). Arrows indicate double-labeled cells. Histograms illustrate the mean (± SD) number of EdU+/PMP2+ oligodendrocytes in central and peripheral regions of the retina. Each dot represents one biological replicate. Significance (*p<0.01) of difference was determined by using a Student’s t-test. Abbreviations: ONL – outer nuclear layer, INL – inner nuclear layer, IPL – inner plexiform layer, GCL – ganglion cell layer, GS – glutamine synthetase. (**j**) scRNA-seq was used to verify patterns of immunolabeling in NIRG cells and oligodendrocytes. These cells were isolated with UMAP ordering maintained for libraries from control and NMDA-damaged retinas. Heatmaps were generated to illustrate patterns of expression of *FABP7, FABP5, PMP2, SOX10, OLIG2* and *ELAVL4*.

### FABPs in eRPC, MGPCs and CMZ progenitors

We next sought to directly compare levels of FABPs between different types of retinal progenitor cells. We aggregated progenitors from E5 and E8 retinas, and MGPCs from retinas treated with NMDA and/or insulin and FGF2 (Fig. 4a). UMAP-ordering of cells revealed 2 distinct clusters of cells for eRPCs and MGPCs (Fig. 4b), with both clusters expressing high levels of proliferation-associated genes including *PCNA, SPC25, TOP2A* and *CDK1* (Fig. 4b,c). MGPCs express high levels of *FABP5* and *PMP2* with significantly higher levels in MGPCs from retinas treated with insulin and FGF2, whereas eRPCs did not express *FABP5* or *PMP2* (Fig. 4c-f). By contrast, levels of *FABP7* were higher in proliferating eRPCs than in proliferating MGPCs with significantly lower levels in MGPCs from retinas treated with insulin and FGF2 (Fig. 4f).

**Figure 4.**
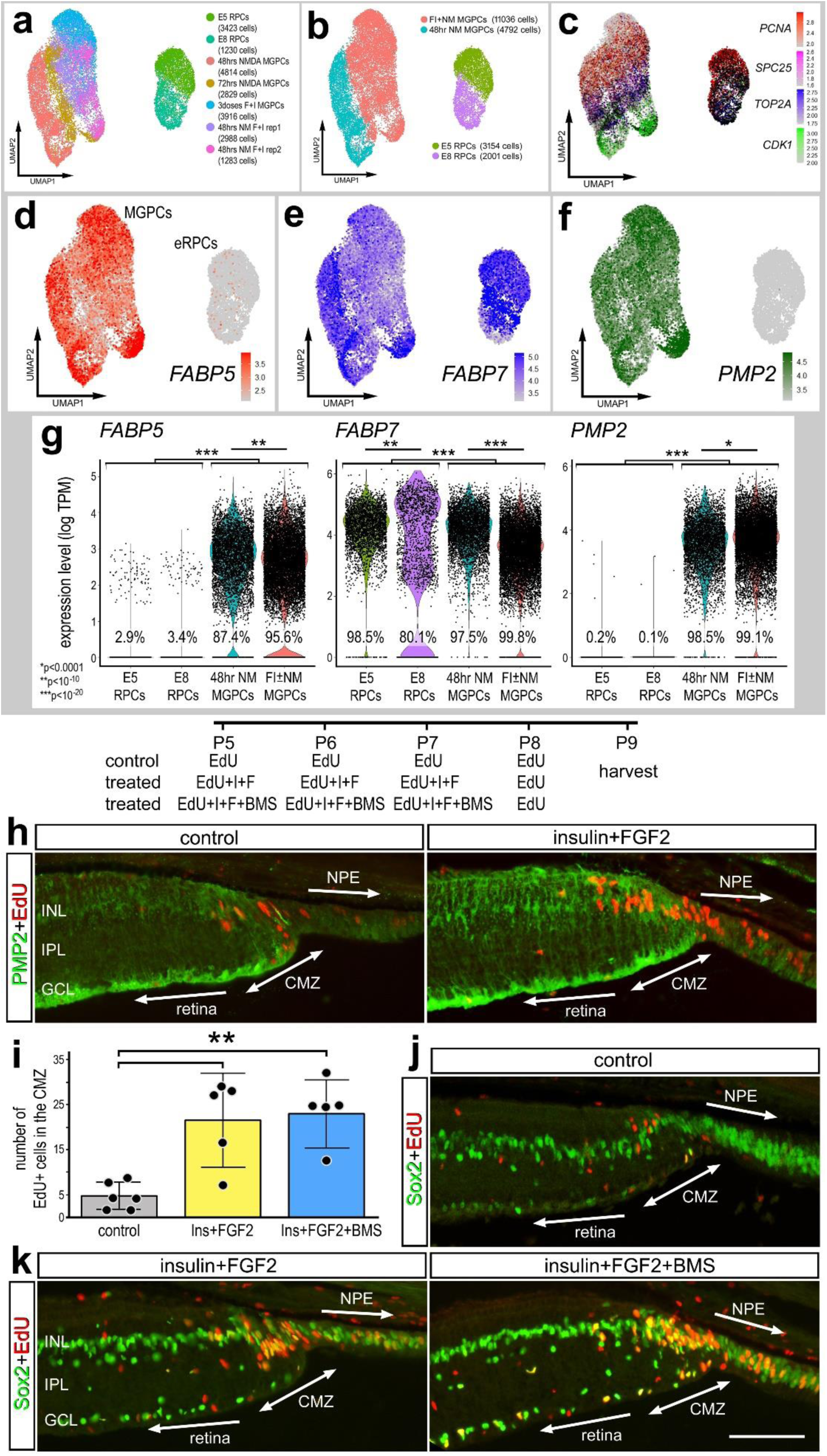
Embryonic progenitors and MGPCs express different FABPs. MG were isolated from scRNA-seq libraries with significant numbers of MGPCs (retinas treated with NMDA and/or FGF2+insulin) and significant numbers of retinal progenitor cells (E5 and E8 retinas). These cells were aggregated and ordered in UMAP plots (**a**-**c**). These unique clusters were probed for levels of expression of *FABP5*, *FABP7*, and *PMP2* in UMAP heatmap plots (**d**-**f**), and quantified in violin plots (**g**). Each dot in violin and scatter plots represents an individual cell. Significant changes (*p<0.1, **p<0.0001, ***p<<0.0001) in levels were determined by using a Wilcox rank sum with Bonferroni correction. The eyes of post-hatch chicks were injected with EdU, insulin +FGF2 and/or FABP inhibitor (BMS309403) (**h**-**k**). Inhibition of FABPs had no effect upon the proliferation of CMZ progenitors (**i**-**k**). Sections of the far peripheral retina and CMZ were labeled for EdU-incorporation (red; **h**,**j,k**,) and antibodies to PMP2 (green; **h**) or Sox2 (green; **j,k**). The histogram in **i** represents the mean (± SD) and each dot represents one biological replicate retina. The calibration bar in **k** represents 50 µm applies to **h**, **j** and **k**. Abbreviations: ONL – outer nuclear layer, INL – inner nuclear layer, IPL – inner plexiform layer, GCL – ganglion cell layer, CMZ – circumferential marginal zone. Significance of difference (**p<0.01) in mean numbers of EdU+ cells in the CMZ and peripheral INL was determined by using ANOVA followed by a post-hoc t-test with Bonferroni correction.

We next examined whether retinal progenitors in the circumferential marginal zone (CMZ) expressed PMP2. Proliferating retinal progenitors with limited neurogenic potential are known to reside in a CMZ in the post-hatch chick eye (Fischer and Reh, 2000). Since our scRNA-seq databases did not include CMZ progenitors at the far peripheral edge of the retina, we immunolabeled sections of CMZ with antibodies to PMP2. PMP2-immunolabeling was present at relatively low levels in MG and CMZ progenitors at the far peripheral edge of the retina (Fig. 4h). Levels of PMP2-immunoreactivity appeared elevated in MG and CMZ progenitors in retinas treated with insulin and FGF2 (Fig. 4h). Although, the proliferation of CMZ progenitors was increased by injections of insulin and FGF2, the proliferation of CMZ progenitors was unaffected by FABP inhibitor (Fig. 4i-k).

### FABPs in retinas treated with insulin and FGF2

We next examined FABP expression in MG and MGPCs in the absence of neuronal damage. In the chick retina the formation of proliferating MGPCs can be induced by consecutive daily injections of Fibroblast growth factor 2 (FGF2) and insulin in the absence of neuronal damage (Fischer et al., 2002; Fischer et al., 2009b; Fischer et al., 2014b; Ritchey et al., 2012). Eyes were treated with two or three consecutive daily doses of FGF2 and insulin, and retinas were processed to generate scRNA-seq libraries. UMAP ordering of cells revealed distinct clusters that were segregated based on cell type (Fig. 5a). MG glia were identified based on the expression of *VIM, GLUL* and *SLC1A3,* and MGPCs were identified based on expression of *TOP2A, NESTIN, CCNB2* and *CDK1* (Fig. 5b). Resting MG from saline-treated retinas formed a cluster of cells distinct from MG from retinas treated with either two or three doses of FGF2+insulin (Fig. 5b). Additionally, MG treated with 2 versus 3 doses of insulin and FGF2 were sufficiently dissimilar to follow different pseudotime trajectories, with MGPCs from retinas treated with 3 doses of insulin+FGF2 occupying a distinct branch (Fig. 5e). Similar to patterns of expression seen in NMDA-damaged retinas, *FABP5, FABP7* and *PMP2* were significantly increased in MG treated with insulin and FGF2 (Fig. 5d). Reduction of pseudotime trajectories to one axis revealed increases in levels of *FABP5* and *PMP2* across pseudotime that paralleled increasing numbers of proliferating MGPCs that expressed *CDK1*, whereas levels of *FABP7* were relatively unchanged across pseudotime (Fig. 5f).

**Fig 5.**
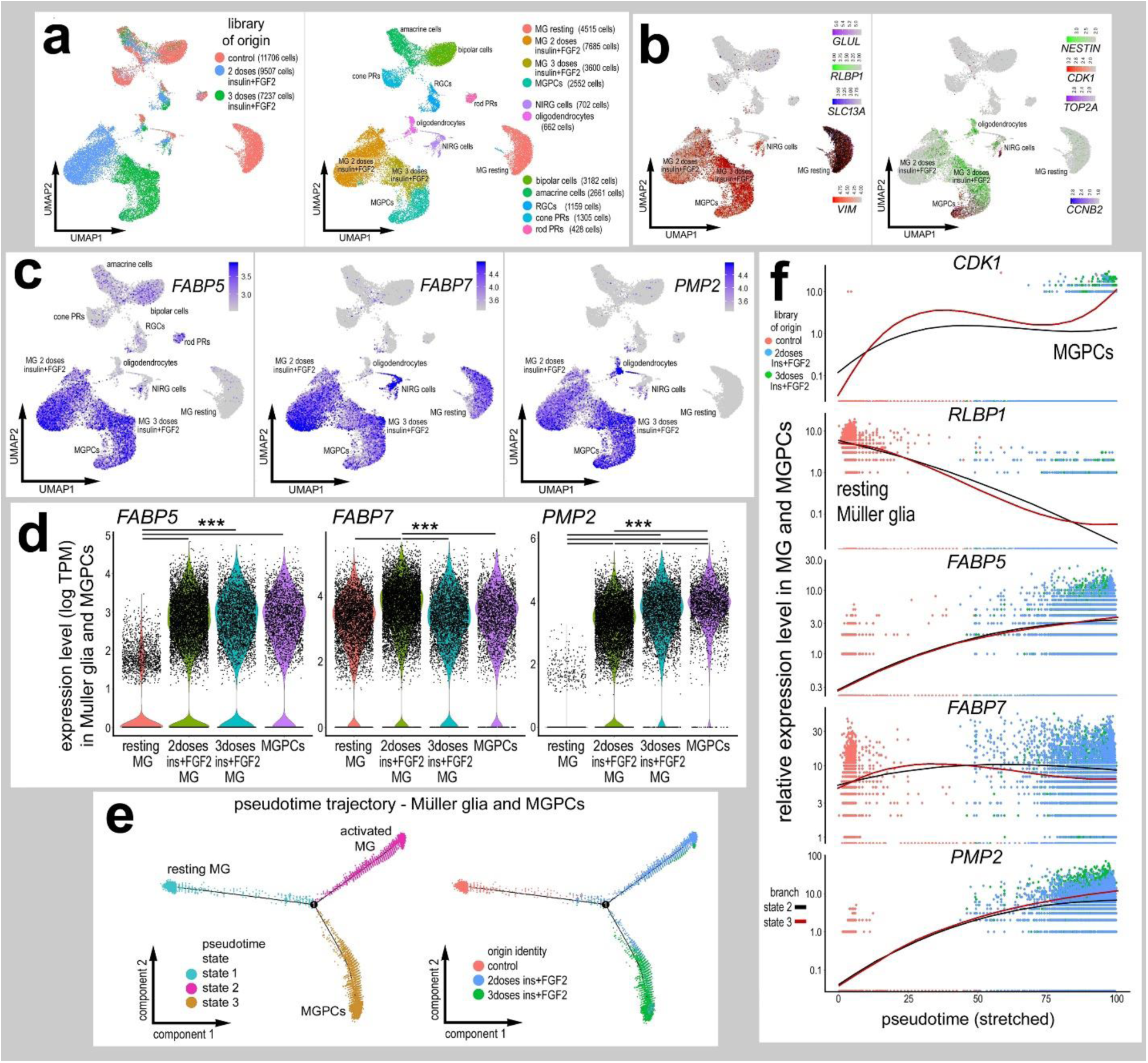
FGF2 and insulin induce expression of PMP2 and FABP5 in MG. scRNA-seq were established for chick retinas treated with saline or two or three doses of FGF2 and insulin (**a**). UMAP ordering of cells revealed distinct clusters of retinal cell types (**a**). Resting MG and growth factor-treated MG formed distinct clusters based on patterns of expression including *GLUL, RLBP1* and *SLC1A3* (**b**). MGPCs were identified based on patterns of expression of *NESTIN, CDK1* and *TOP2A* (**b**). UMAP heatmaps and violin plots were established to illustrate patterns and levels of expression for *FABP5, FABP7* and *PMP2* (**c** and **d**). MG were re-embedded and ordered across pseudotime with a junction and trajectories populated by activated MG (mostly 2 doses of insulin and FGF2) or proliferating MGPGs branches (mostly 3 doses of insulin and FGF2) (**e**). Dimensional reduction of pseudotime to the X-axis placed resting MG (high *RLBP1*) to the left and activated MG (black line) and MGPCs (red line; high *CDK1*) to the right (**f**). Levels of *FABP5, FABP7* and *PMP2* were plotted across pseudotime (**f**).

We next compared levels of FABPs in MG and MGPCs in retinas treated with insulin+FGF2 ± NMDA. We probed a large aggregate scRNA-seq library of more than 55,000 cells wherein UMAP ordering revealed discrete clustering of resting MG, activated MG and MGPCs (supplemental Fig. 2a). Levels of *FABP5* and *PMP2* were lowest in resting MG, and highest in activated MG at 24hrs after NMDA and MGPCs compared to the elevated levels seen in MG treated with insulin+FGF2, NMDA+insulin+FGF2 and activated MG at 48+72 hrs after NMDA (supplemental Fig. 2f-h). Patterns of expression for *FABP7* were similar to those seen for *FABP5* and *PMP2,* although higher levels of expression were observed in both reactive and, in particular, resting MG (supplemental Fig. 2f-h). By comparison, levels of *FASN* were highest in resting MG, lowest in MGPCs, and at intermediate levels in activated MG observed following injury or growth factor treatment (supplemental Fig. 2f-h).

To test whether PMP2 was increased when MG were treated with FGF2 we immunolabeled retinal sections. We found that PMP2-immunoreactivity was increased in FGF2-treated MG (Fig. 6a). This effect was not evident in central regions of the retina, but was apparent in peripheral regions (Fig. 6a). Some of the PMP2+/Sox2+ MG were labeled for EdU (Fig. 6b), indicating that these cells were proliferating MGPCs. We next investigated whether FABP-inhibition influenced the formation of MGPCs in the absence of neuronal damage. We have reported previously that proliferating MGPCs are formed in peripheral regions of retinas treated with consecutive daily injections of FGF2 alone (Fischer et al., 2014). Accordingly, we applied BMS309403 with FGF2 and probed for the formation of proliferating MGPCs. Inhibition of FABPs significantly reduced numbers of Sox2/EdU-labeled cells in the INL of FGF2-treated retinas (Fig. 6c,d).

**Figure 6.**
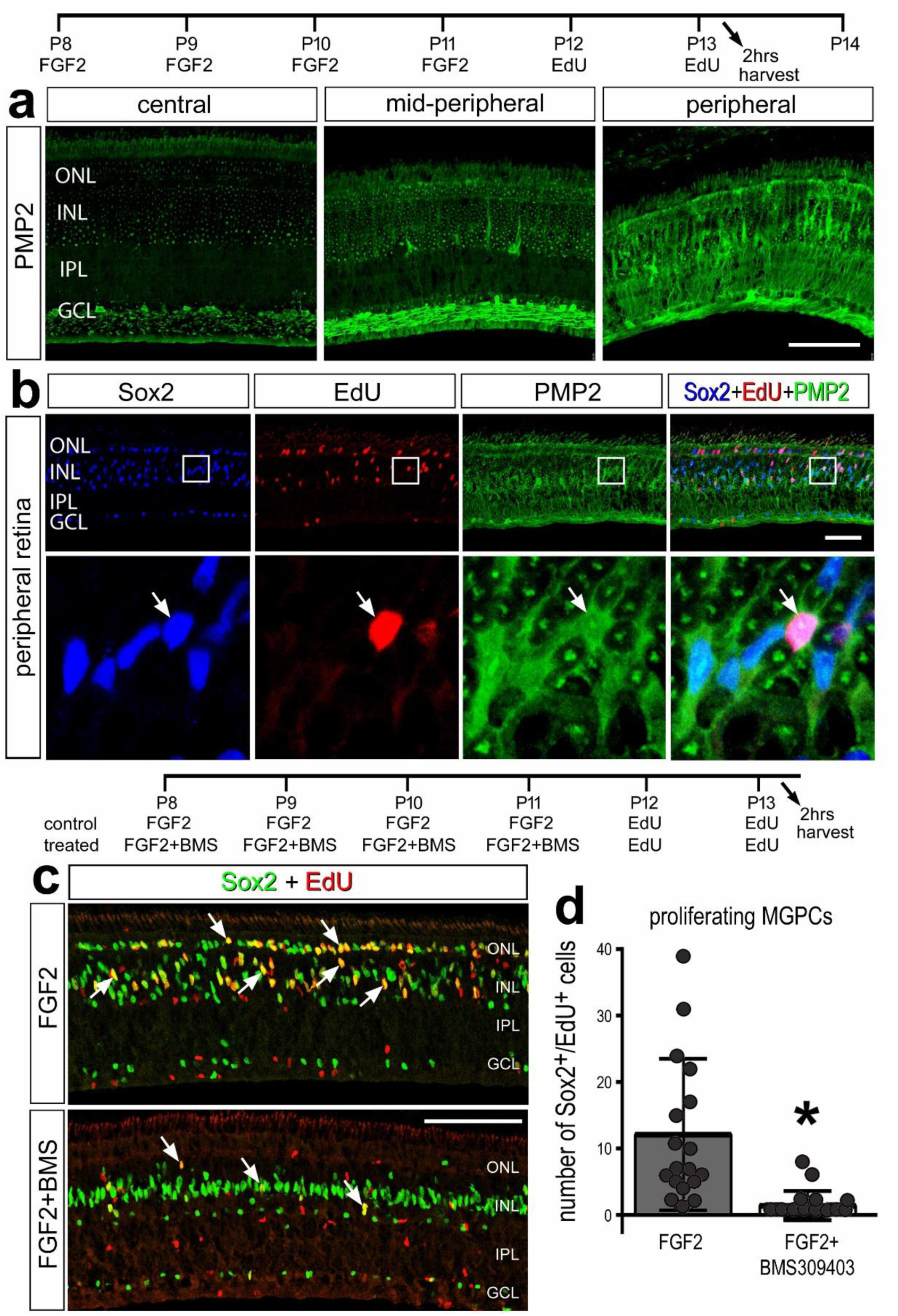
Treatment of retinas with FGF2 stimulates MG to up-regulate PMP2 and proliferate, and this proliferation can be blocked by FABP-inhibitor. (**a,b**) Eyes were treated with 4 consecutive daily intraocular injections of FGF2, followed by 2 consecutive daily injections of EdU, and retinas harvested 2hrs after the last injection. (**c,d**) Alternatively, eyes received 4 consecutive daily injections of FGF2 ± BMS309403, followed by 2 consecutive daily injections of EdU, and eyes harvested 24hrs after the last injection. Sections of the retina were labeled for EdU (red; **b**) and antibodies to PMP2 (green; **a,b**) or Sox2 (blue in **b**, green in **c**). Arrows indicate MG nuclei labeled for EdU and Sox2. The histogram in **d** represents the mean (± SD) and each dot represents one biological replicate retina. Significance (*p<0.01) of difference was determined by using a Student’s t-test. The calibration bar represents 50 µm.

### Genes regulated by FABP-inhibition

To identify transcriptional changes downstream of FABP inhibition we generated scRNA-seq libraries for control retinas and retinas treated with inhibitor (BMS309403) ± NMDA. Aggregation of the scRNA-seq libraries revealed distinct UMAP clusters of retinal neurons and glia (Fig. 7a,b). UMAP-ordering did not result in distinct clustering of neurons based on treatment, whereas MG were clustered according to treatments (Fig. 7a,b). Genes upregulated in activated MG included *HBEGF, TUBB6* and *TGFB2* (Fig. 7d). MGPCs upregulated genes such as *CDK1, TOP2A* and *ESPL1* (Fig. 7e). Resting MG expressed high levels of *GLUL, RLBP1* and *SLC13A* which were downregulated in activated MG (Fig. 7f). We identified differentially expressed genes (DEGs) in MG from retinas treated with BMS, NMDA and BMS+NMDA (Fig. 7c; supplemental tables 1,2,3). BMS treatment of undamaged retinas resulted in significant changes in gene expression, which included down-regulation of 393 genes including markers for resting mature glia such as *GLUL, CRABP-I, RLBP1, CA2, SFRP1* and *ID4* (Fig. 7g, supplemental table 1). By comparison, BMS-treatment resulted in an up-regulation of 495 genes including FABPs, secreted factors associated with glial activation and receptors for TNF-related ligands (Fig. 7g, supplemental table 1). Gene ontology (GO) enrichment analysis of DEGs from saline-BMS-treated MG revealed significant enrichment for up-regulated genes associated with gliogenesis, wound healing and developmental processes (Fig. 7i), whereas down-regulated genes were associated with neurogenesis and cell proliferation (Fig. 7i). BMS-treatment of NMDA-damaged retinas resulted in significant changes in gene expression which included down-regulation of 192 genes including markers for proliferation, pro-glial genes *NFIX* and *ID4*, and High Mobility Group (HMG) genes (Fig. 7h, supplemental table 2). By comparison, BMS-treatment resulted in an up-regulation of 114 genes including *FABP5* and *FABP7,* secreted factors *BMP4, WNT4* and *WNT6*, and glial reactivity genes such as *CD44* (Fig. 7h, supplemental table 2). GO enrichment analysis of down-regulated genes from NMDA/BMS-treated MG revealed significant enrichment for genes associated with proliferation and organelle fission (Fig. 7i). By comparison, GO enrichment analysis for upregulated genes from NMDA/BMS-treated MG revealed significant enrichment for genes associated with growth factor signaling, cell death and cell motility (Fig. 7j).

**Figure 7.**
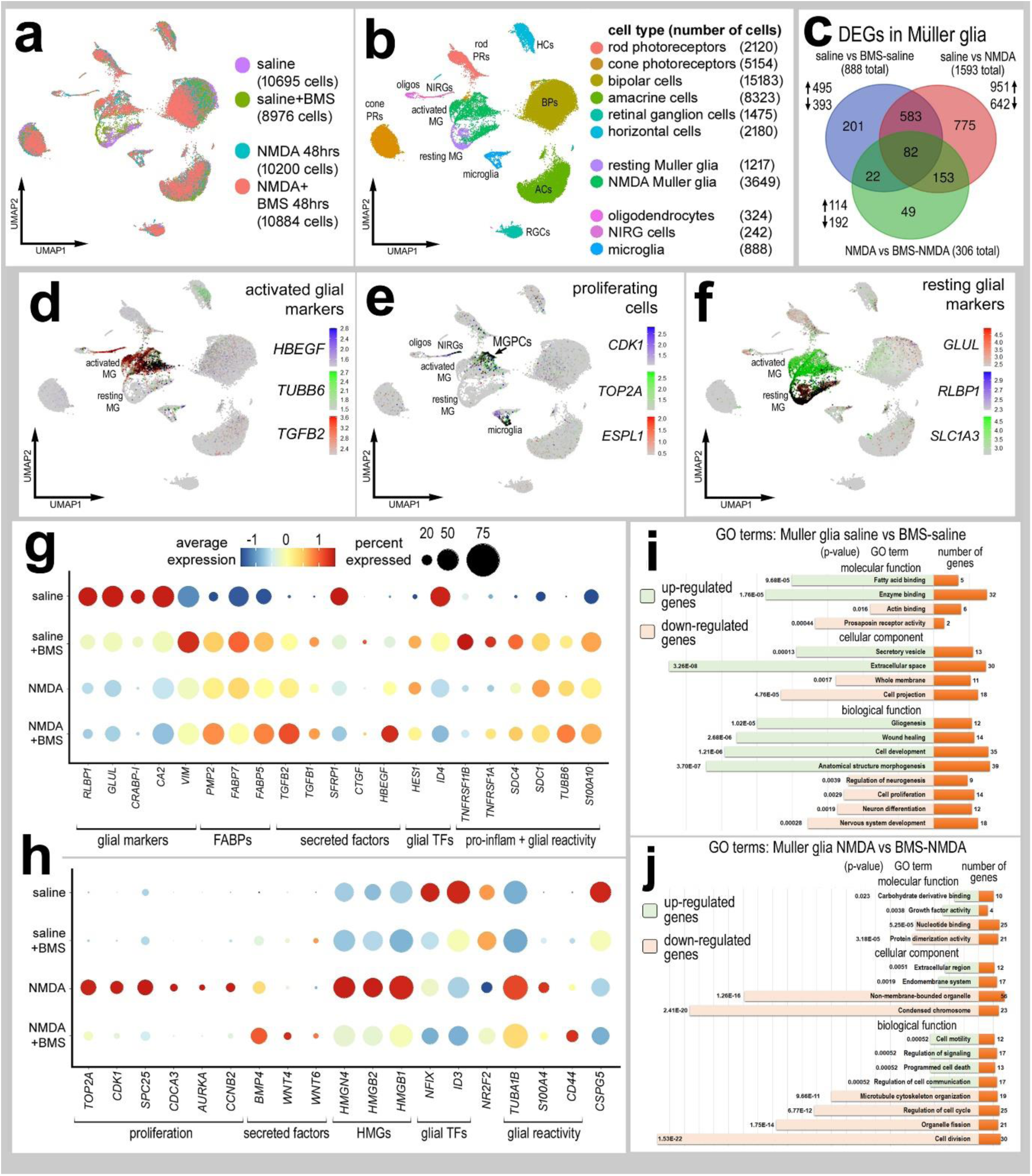
Inhibition of FABP significantly impacts single cell transcriptomic profiles of MG. Retinas were treated with saline ± BMS309403 or NMDA ± BMS309403, and scRNA seq libraries were generated to analyze changes in MG gene expression. UMAP ordering of cells was established and MG were identified based on expression of genes associated with activated glia (**d**), proliferating MGPCs (**e**), and resting glia (**f**). Differentially expressed genes (DEGs) were identified for MG from retinas treated with saline vs BMS-saline, saline vs NMDA, and NMDA vs BMS-NMDA and plotted in a Venn diagram (**c**). Dot plots indicating the percentage of expressing MG (size) and expression levels (heatmap) for genes related to resting glia, secreted factors, glial transcription factors, inflammation, glial reactivity and proliferation (**g, h**). All genes displayed in the Dot plot have significantly different (p<0.0001) expression levels in MG from retinas treated with saline vs saline-BMS (**g**) or in MG from retinas treated with NMDA vs NMDA-BMS (**h**). Gene Ontology (GO) terms for the enriched genes in the BMS treated and BMS+NMDA treated were compiled (ShinyGO) and grouped by biological process, cellular component and molecular function. GO enrichment analysis was performed. The significance of the function and the number of enriched genes are listed for each GO category.

We isolated MG from 4 treatment groups and re-ordered these cells in UMAP plots. The MG formed 5 distinct clusters; 2 clusters of resting MG (occupied predominantly by cells from retinas treated with saline- and BMS-saline), 2 clusters of activated glia (occupied predominantly cells from retinas treated with NMDA, BMS-NMDA and BMS-saline), and MGPCs (occupied predominantly by cells from retinas treated with NMDA alone) (supplemental Fig. 2a-c). This finding suggest that BMS-treatment induces MG to acquire a reactive phenotype. Markers for mature resting MG were significantly down-regulated in activated MG and MGPCs; these markers included *GLUL, RLBP1, CSPG5* and *ID4* (supplemental Fig. 2d-h). By contrast, activated MG significantly up-regulated markers associated with reactivity such as *HBEGF, TGFB2, BMP4, S100A, SDC4, CD44, FABP5, FABP7* and *PMP2* (supplemental Fig. 2d-h). Not surprising, MGPCs had elevated levels of proliferation markers such as CDK1, TOP2A and SPC25 (supplemental Fig. 2d-h). GO enrichment analysis for DEGs between resting MG and activated MG revealed significant enrichment for up-regulated genes associated with cell metabolism, structural cytoskeleton, and organelle organization (supplemental Fig. 2j). By comparison there was significant enrichment for down-regulated genes associated with transcriptional regulation, cell adhesion, cellular projections, and neuronal development (supplemental Fig. 2j). GO enrichment analysis for DEGs between activated MG and MGPCs revealed significant enrichment for up-regulated genes associated with cell division, and enrichment for down-regulated genes associated with nervous system development and neuronal differentiation (supplemental Fig. 2i).

We next isolated the MG, microglia and NIRG cells, re-embedded these cells in UMAP plots and probed for cell signaling networks and putative ligand-receptor interactions using SingleCellSignalR (Cabello-Aguilar et al., 2020). We chose to focus our analyses on the MG, microglia and NIRG cells because there is significant evidence to indicate autocrine and paracrine signaling among MG, microglia and NIRG cells in the context of glial reactivity and the formation of proliferating MGPCs (Fischer et al., 2014b; Wan et al., 2012; White et al., 2017; Zelinka et al., 2012a). Resting MG included cells for saline and BMS-treatment groups, activated MG included cells mostly from BMS-saline, NMDA and BMS-NMDA treatment groups, and MGPCs were predominantly derived the NMDA-treatment group (Fig. 8a-c). Numbers of LR-interactions (significant upregulation of putative ligand and receptor) between cell types in the different treatment groups varied between 70 and 315 (Fig. 8d-g). We performed analyses on glial cells from each treatment group and compared changes across the most significant LR-interactions. For example, LR-interactions included *IGF1R* and *FGFR1* in activated MG to MGPCs, but not when treated with BMS (Fig. 8f,g). By comparison, BMS-treated MG included LR-interactions with *TGFBR2* and interactions involving *TNFRSF11B*, *MDK, PTN* and *PTPRZ1* (Fig. 8f,g). We compared significant changes in LR-interactions among glial cells and interactions unique to treatment groups for undamaged and damaged retinas. We identified 33 LR-interactions specific to saline-treated glia and 148 LR-interactions specific to BMS-treated glia in undamaged retinas (Fig. 8h,i,l,m,p). Glia in undamaged saline-treated retinas included LR-interactions for *FGF9-FGFR3/4*, *BMP2-ACVR2* and *BMP6-BMPR2* (Fig. 8h,i,l,m,p). By comparison, glia in undamaged BMS-treated retinas included LR-interactions associated with activated glial phenotypes such as *IL1B-IL1RAP*, *TGFB1/2-TGFBR2*, *HBEGF-CD9/CD82/ERBB2/EGFR*, and *JAG1/JAG2/PSEN1-NOTCH2* (Fig. 8h,i,l,m,p p). We identified 40 LR-interactions specific to saline/NMDA-treated glia and 86 LR-interactions specific to BMS/NMDA-treated glia in damaged retinas (Fig. 8j,k,n,o,q). LR-interactions unique to glia in NMDA-damaged retinas included *BMP*-, *MDK*-, *FGF*- and *DLL1-Notch1*-signaling (Fig. 8j,k,n,o,q). By comparison, LR-interactions unique to BMS/NMDA-damaged retinas included *JAG1/2/PSEN1-Notch2*, *INHBA-ACVR2*, *TGFB1-ITGB3* and *WNT5A-LRP2/FZD4* (Fig. 8 j,k,n,o,q).

**Figure 8.**
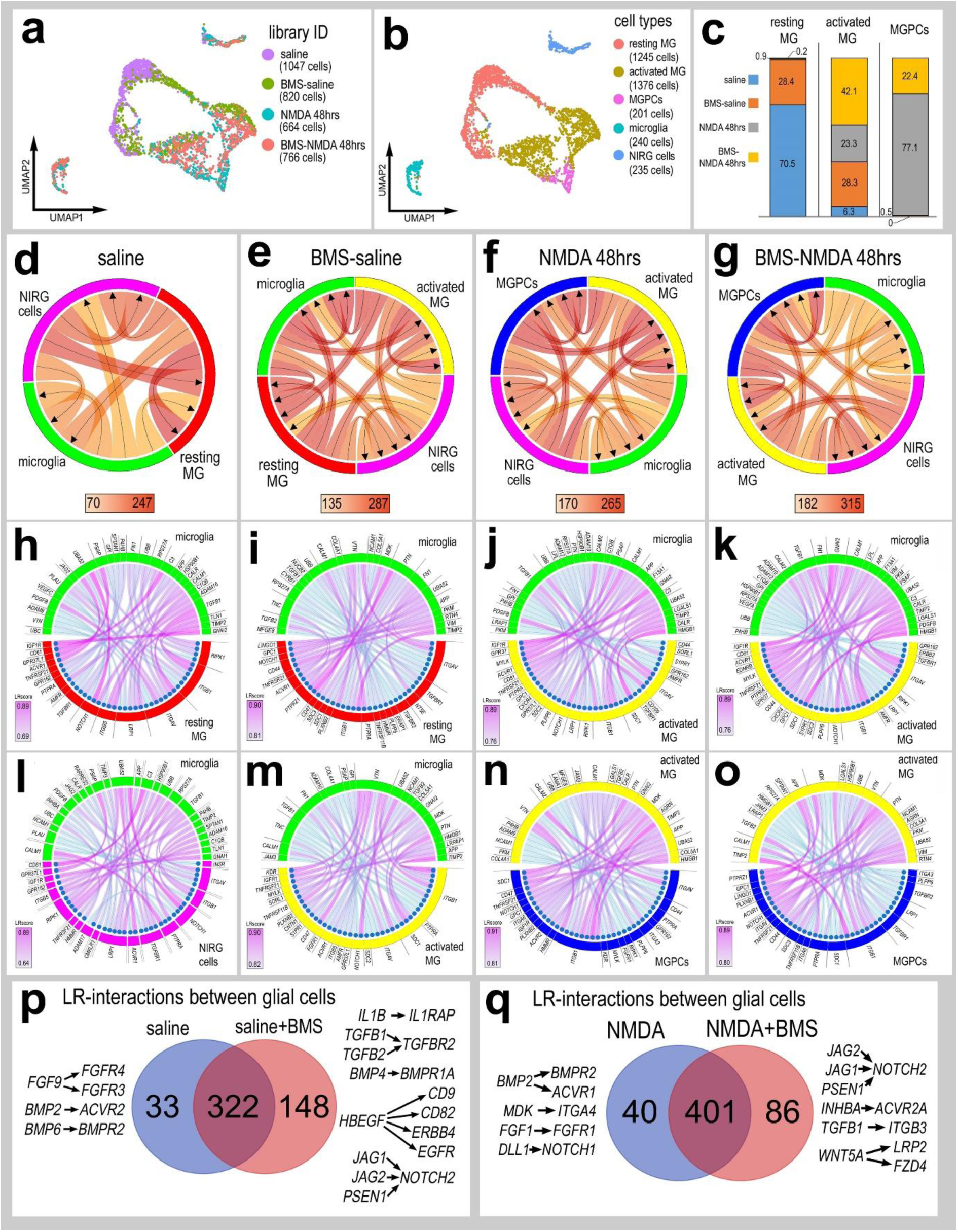
Ligand-receptor (LR) interactions inferred from scRNA-seq data between microglia, NIRG cells, resting MG, activated MG and MGPCs. Retinal glia were isolated, re-embedded and ordered in UMAP plots to reveal distinct clusters of cells (**a**, **b**). The resting MG were comprised primarily of MG from saline-treated retinas, activated MG were comprised of cells from BMS-saline, NMDA and BMS-NMDA treated retinas, and MGPCs were primarily comprised of cells from NMDA-treated retinas (**a**-**c**). Glia from different treatment groups were analyzed using SingleCellSignalR to generate chord diagrams and illustrate potential autocrine and paracrine LR-interactions (**d**-**g**). LR-interactions were identified for glial cells for different treatment groups including saline (**h,l**), BMS-saline (**i,m**), NMDA (**j,n**) and BMS-NMDA (**k,o**). For each treatment group, the 40 most significant LR-interactions between microglia to resting MG (**h,i**), microglia to NIRG cells (**l**), microglia to activated MG (**j,k,m)**, and activated MG to MGPCs (**n-o**) were identified and illustrated in chord plots and LRscore heat maps. Treatment-specific differences in glial LR-interactions in saline vs BMS-saline (**p**) and NMDA vs BMS-NMDA (**q**) are illustrated in Venn diagrams with some key interactions listed.

### Inhibition of FABPs in resting MG

We next sought to investigate and validate changes in cell signaling in damaged retinas treated with FABP-inhibitor. One day after treatment with NMDA ± BMS there was a significant increase in numbers of dying TUNEL+ cells (Fig. 9a,b). By contrast, we observed a significant decrease in pS6 in MG in damaged retinas treated with BMS (Fig. 9a,b), consistent with findings from SingleCellSignalR where LR-interactions for *FGF1-FGFR1* and *MDK-ITGA4* are missing with BMS-treatment (Fig. 8q). MDK and FGF are known to active mTor-signaling and up-regulated pS6 in MG in the chick retina (Campbell et al., 2021a; Zelinka et al., 2016). Consistent with these findings, we observed reduced levels of pStat3 in MG nuclei in damaged retinas treated with BMS (Fig. 9a,b). Stat phosphorylation is known to be down-stream of PDGF-signaling (Li et al., 2012; Popielarczyk et al., 2019) and PDGFA-PDGFRA LR-interactions are missing from damaged retinas treated with BMS (Fig. 8q). Similarly, we find reduced levels of pSMAD1/5/8 in MG nuclei in damaged retinas treated with BMS (Fig. 9a,b). This may result from increased signaling through ACVR2A and TGFB1/ITGB3 which may antagonize BMP/SMAD-signaling (Todd et al., 2017).

**Figure 9.**
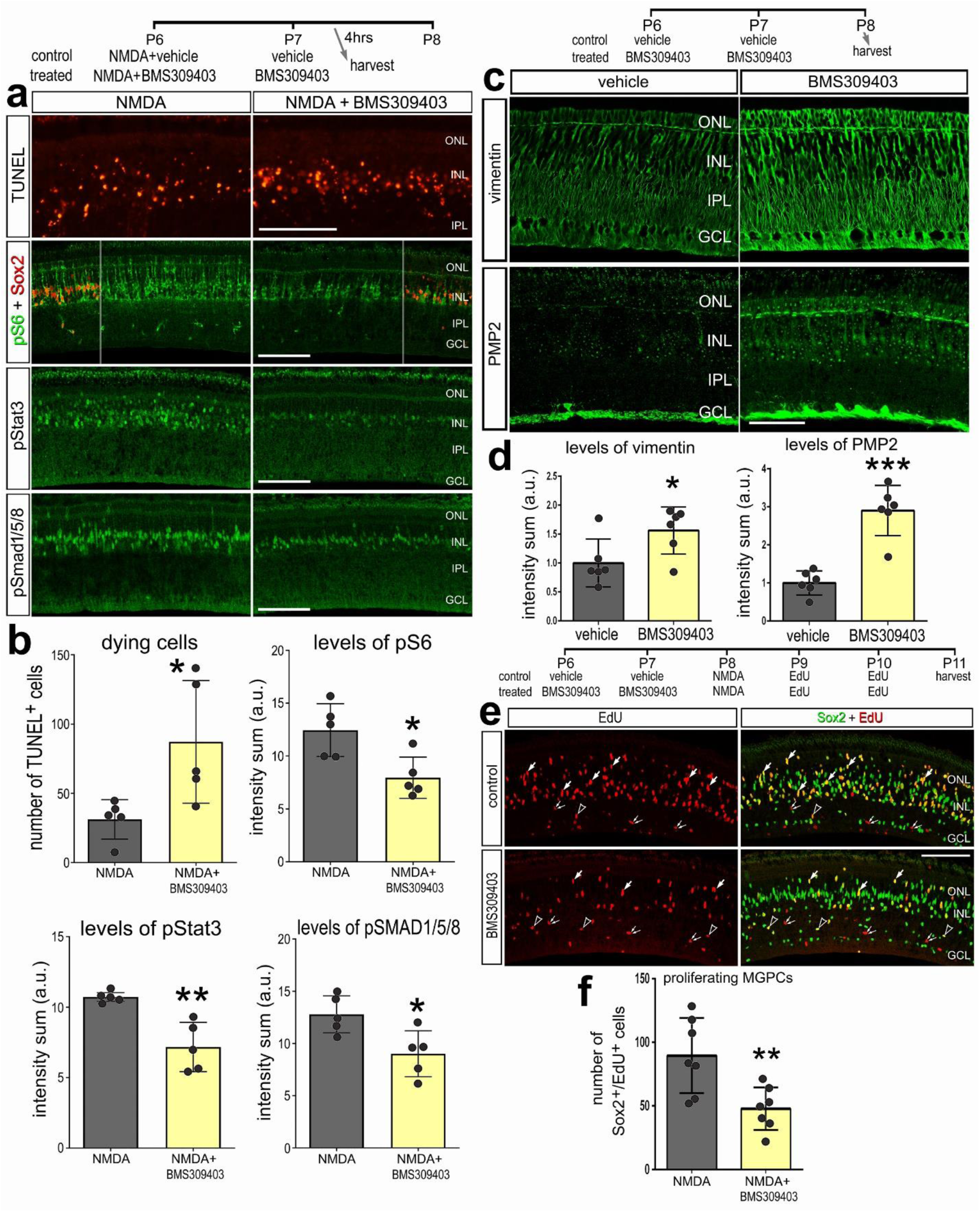
(**a**-**b**) Inhibition of FABPs suppresses multiple cell signaling pathways in NMDA-damaged retinas. BMS309403 or vehicle were injected at P6 with NMDA, followed by injection of vehicle or BMS309403 at P7, and retinas were harvested 4 hours after the last injection. (**a**) Retinal sections were labeled for fragmented DNA (TUNEL; red) or antibodies to pS6 (green) and Sox2 (red), pStat3 (green), or pSmad1/5/8 (green). The histograms in **b** illustrate numbers of dying cells or immunofluorescence intensity sum for pS6, pStat3, and pSmad1/5/8. (**c**-**d**) Inhibition of FABPs in undamaged retinas up-regulates vimentin and PMP2 in MG, and suppresses the formation of MGPCs when damage is induced. Sections of the retina were labeled with antibodies to vimentin or PMP2 (**c**). The histogram in **d** illustrates levels of immunofluorescence intensity sum for vimentin and PMP2. (**e**-**f**) BMS309403 was injected at P6 and P7, NMDA was injected at P8, EdU was injected at P9 and P10, and retinas were harvested at P11. Retinal sections were labeled for EdU and antibodies to Sox2 (**e**). Arrows indicate the nuclei of MG labeled for Sox2 and EdU in the INL, hollow arrow-heads indicate the nuclei of NIRG cells labeled for Sox2 and EdU in the IPL, and small double-arrows indicate putative proliferating microglia labeled for EdU alone. The calibration bars in **a, c** and **e** represents 50 µm. The histogram in **f** illustrates numbers of Sox2+/EdU+ MGPCs. Abbreviations: ONL – outer nuclear layer, INL – inner nuclear layer, IPL – inner plexiform layer, GCL – ganglion cell layer. The histograms illustrated the mean (± SD) and each dot represents one biological replicate retina. Significance (*p<0.05, **p<0.01, ***p<0.001) of difference was determined by using a Student’s t-test.

scRNA-seq data indicated that BMS-treatment of undamaged retina had a significant impact on the transcriptional profile of MG with profiles resembling de-differentiation and reactive glia. We sought to verify some of the scRNA-seq data by labeling BMS-treated retinas with antibodies to vimentin or PMP2. We found that BMS-treatment significantly increase immunofluorescence for vimentin and PMP2 in MG (Fig. 9c,d). Since, BMS treatment appeared to stimulate MG to become reactive and de-differentiate processes that occur during a transition to a progenitor-like phenotype (Hoang et al., 2020), we tested whether BMS treatment primes MG to become proliferating progenitors in damaged retinas. BMS treatment should have primarily inhibited FABP7 in resting MG and this inhibition was predicted to enhance the ability of MG to become proliferating MGPCs. Contrary to expectations, we found that BMS treatment of retinas before NMDA-induced damage suppressed the formation of proliferating MGPCs (Fig. 9e,f). The proliferation of NIRG cells was not affected by BMS treatment prior to NMDA-induced retinal damage (not shown).

### Effects of FABP-inhibition in microglia

Retinal microglia and infiltrating macrophage are known to promote the formation of MGPCs in chick and zebrafish retinas (Fischer et al., 2014b; Palazzo et al., 2020; White et al., 2017) and suppress the neuronal differentiation of reprogrammed MG in mouse retinas (Todd et al., 2020). Accordingly, we investigated whether microglia were influenced by treatment with FABP inhibitor. The application of BMS to NMDA-damaged retinas suppressed the accumulation of microglia and significantly reduced numbers of EdU+/CD45+ cells (Fig. 10a,b). The microglia in BMS-NMDA treated retinas appeared to retain a reactive morphology (Fig. 10a). Thus, it is possible that reduced numbers of proliferating MGPCs resulted, in part, from reduced accumulation of reactive monocytes in BMS-treated retinas.

**Figure 10.**
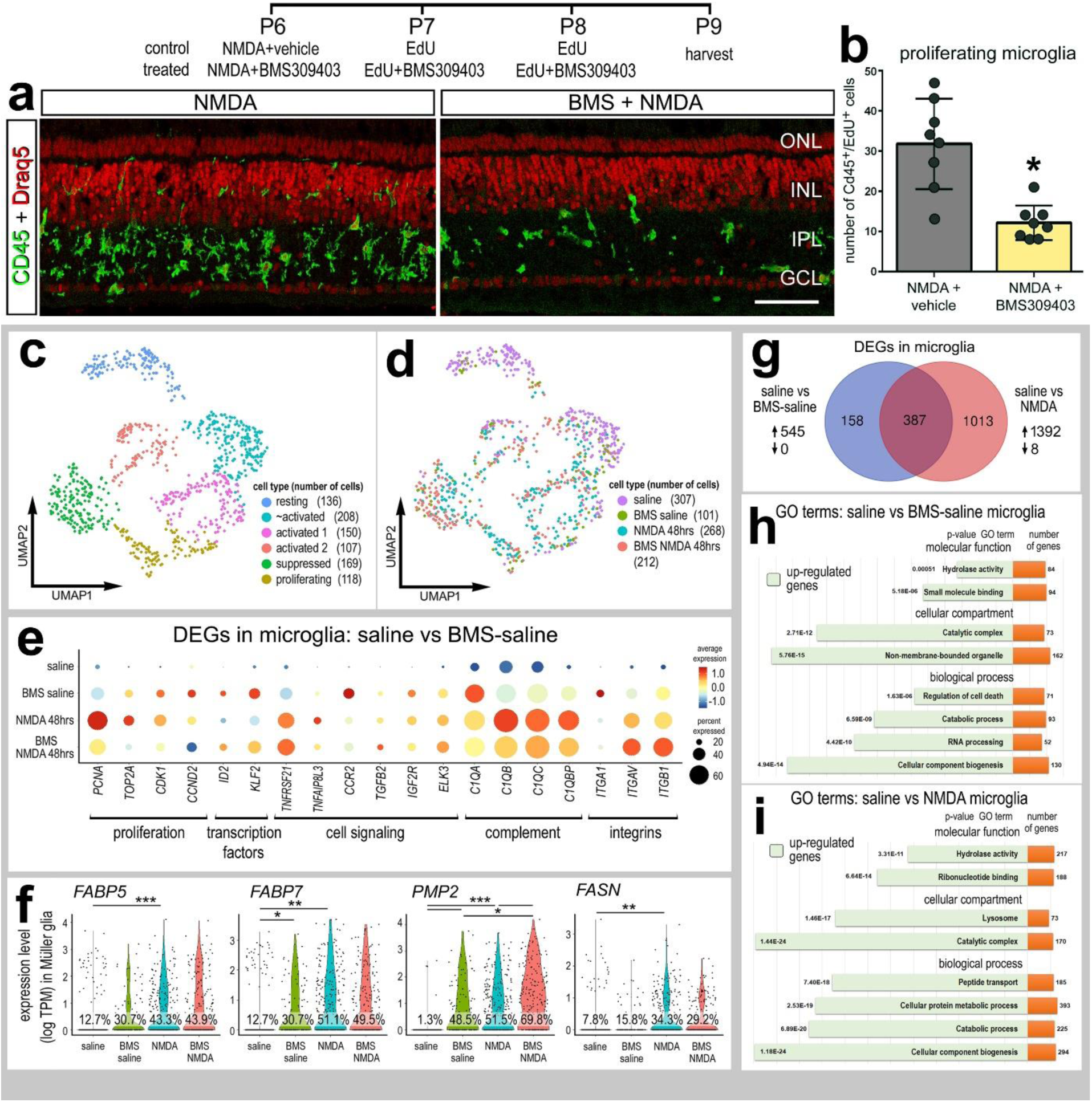
FABP inhibition reduces microglia proliferation. Inhibition of FABPs with BMS309403 suppressed the accumulation and proliferation of microglia (**a**,**b**). Sections of the retina were labeled for DRAQ5 (red) and CD45 (green; **a**). The calibration bar in **a** represents 50 µm. Abbreviations: ONL – outer nuclear layer, INL – inner nuclear layer, IPL – inner plexiform layer, GCL – ganglion cell layer. The histogram in **b** represents the mean (± SD) number of CD45+/EdU+ cells and each dot represents one biological replicate. Significance of difference (*p<0.01) was determined using a two-way paired t-test. scRNA-seq libraries established from retinas treated with saline ± BMS or NMDA ± BMS and microglia were isolated and analyzed for changes in genes expression. The microglia were clustered by UMAP and analyzed for differentially expressed genes (DEGs) which are illustrated in a Venn diagram (**g**). Dot plots indicating the percentage of expressing microglia (size) and expression level (heatmap) were generated for classes of genes including those important for proliferation, transcriptional regulation, cell signaling and inflammatory signaling (**e**). All genes displayed in the dot plot have significantly different (p<0.0001) expression levels in microglia from retinas treated with saline vs saline-BMS. Gene Ontology terms for the enriched genes in the BMS treated and BMS+NMDA treated were compiled (ShinyGO) (**g,h,i**). GO was performed for significantly up-regulated genes (green) and the number of enriched genes in each GO category (orange) are displayed. There were insufficient numbers of down-regulated genes and insufficient DEGs identified for NMDA vs NMDA+BMS to perform GO enrichment analyses. Violin plots of *FABP5, FABP7, PMP2* and *FASN* illustrate the percentage of expressing cells and significant changes in levels of expression between treatment groups (**f**). Significance of difference (*p<0.01, **p<0.001, ***p<<0.001) was determined by using a Wilcoxon rank sum with Bonferroni correction.

We isolated and UMAP-embedded microglia from retinas treated with saline, BMS and NMDA. UMAP plots revealed distinct clusters of resting microglia, activated and proliferating cells (Fig. 10c). The resting microglia UMAP clusters were predominantly occupied by microglia from saline-treated retinas, whereas microglia from BMS-saline treated retinas were clustered among activated cells (Fig. 10c,d). BMS-treatment of microglia in undamaged retinas resulted in up-regulation of 545 genes, whereas no genes were significantly down-regulated (Fig. 10e,g; supplemental table 4). BMS-treatment in undamaged retinas stimulated microglia to up-regulate genes associate with proliferation, cell signaling, complement, integrins and glial transcription factors (Fig. 10e; supplemental table 4). NMDA-treatment stimulated microglia to significantly up-regulate nearly 1400 genes including *FABP5, FABP7, PMP2* and *FASN,* whereas only 8 genes were down-regulated (Fig. 10g,f; supplemental table 5). BMS-treatment of NMDA-damaged retinas resulting if changes in expression of only 6 genes, and *PMP2* was among the few genes upregulated by microglia in BMS/NMDA-damaged retinas (Fig. 10f). GO enrichment analysis of DEGs in microglia from retinas treated with saline ± BMS indicated groups of genes associated with cellular biogenesis, regulation of cell death and hydrolytic/catabolic processes (Fig. 10h). Thus, it is not surprising that the BMS-treated microglia were embedded among microglia from NMDA-damaged retinas in UMAP plots (Fig. 10c,d). GO enrichment analysis of DEGs in microglia from retinas treated with saline ± NMDA indicated groups of genes associated with lytic enzyme activity, lysosomal activity and cellular biogenesis (Fig. 10j). There were only 6 DEGs in microglia from retinas treated with NMDA ± BMS, consistent with the co-clustering of microglia from these treatments in UMAP plots (Fig. 10d). These microglia were harvested at 48hrs after NMDA-treatment and, thus, it is likely that significant differences in gene expression that led to decreased accumulation of reactive microglia in BMS/NMDA-treated retinas occurred shortly after damage and BMS-treatment. This is consistent with scRNA-seq findings in NMDA-damaged mouse retinas wherein microglia significantly up-regulated genes for pro-inflammatory cytokines between 3 and 12 hrs after damage (Todd et al., 2019).

### FASN influences MGPCs, neuronal survival and the accumulation of reactive microglia

Fatty acid synthase (FASN) –dependent fatty acid synthesis is necessary for FABP activity. scRNA-seq data indicated that FASN is widely expressed by most retinal cell types (Fig. 11a). In MG, levels of FASN were elevated in resting MG and down-regulated in MG at 24, 48 and 72 hrs after NMDA treatment, and remained low and prevalent in MGPCs (Fig. 11a,b). Relative levels of FASN in MG and MGPCs across different treatments were highest in resting MG, lowest in MGPCs and intermediate levels in activated MG at different times after NMDA and doses of insulin + FGF2 (supplemental Fig. 2f-h). We further assessed patterns of expression of FASN in aggregate libraries from time-points soon after NMDA, at 3 and 12 hrs after treatment. Levels of *FASN* were significantly up-regulated in MG at 3hrs after NMDA and back down at 12 and 48 hrs after NMDA (Fig. 11c,d), suggesting a rapid and transient need for elevated fatty acid synthesis in MG shortly after NMDA-treatment.

**Figure 11.**
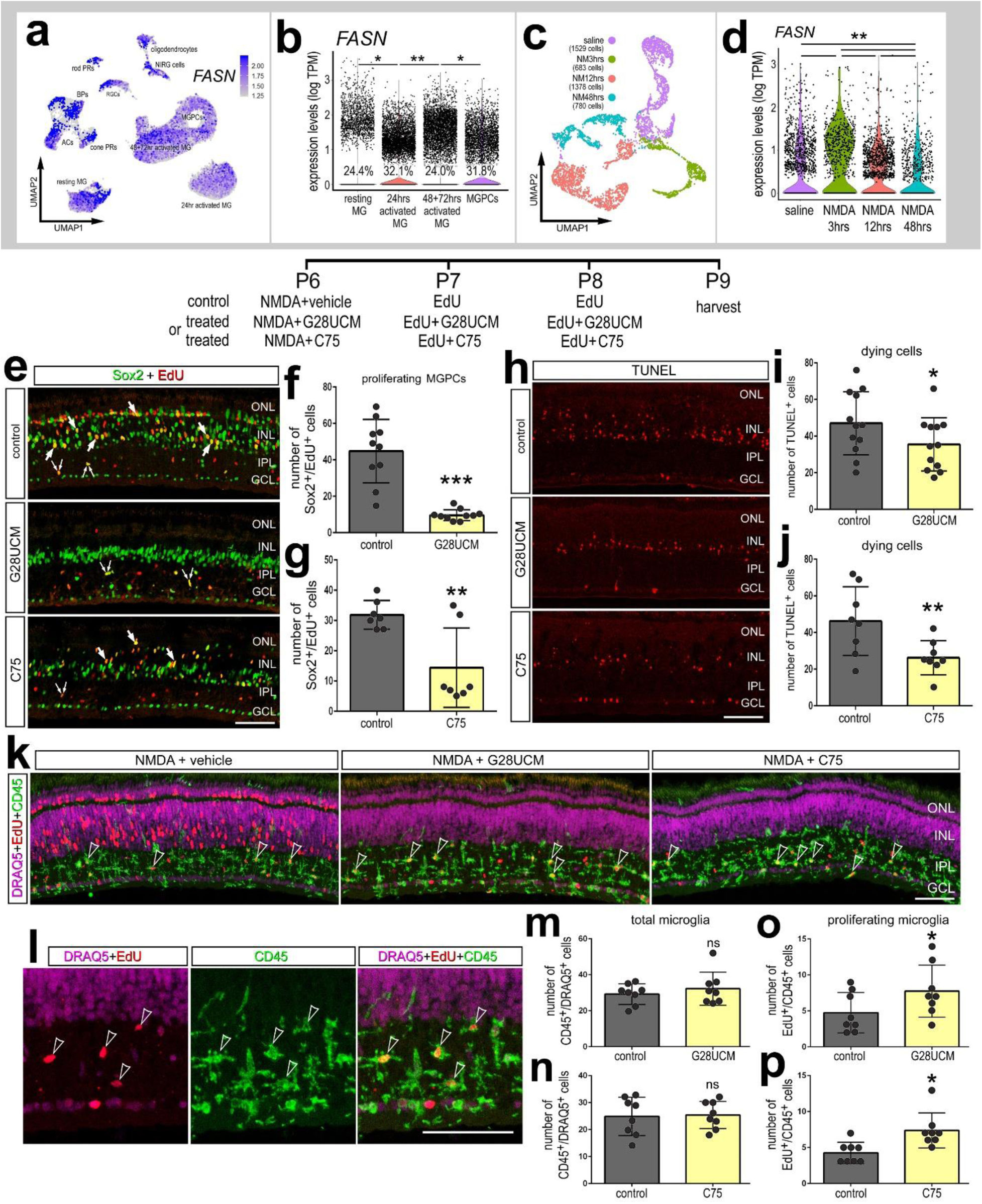
Fatty acids synthase inhibitors significantly reduce MGPC formation. scRNA-seq libraries of the NMDA damaged retinas were probed Fatty acids synthase (FASN). UMAP heatmap plot in **a** illustrate FASN expression across different cells types from retinas treated with saline or NMDA at 24, 48 or 72 hrs after treatment. Expression levels and percentage expressing cells in MG are illustrated in violin plots (**b**,**d**). The UMAP plot in **c** illustrates discrete clustering of MG from retinas treated with saline or NMDA at 3, 12 or 48 hrs after treatment. FASN inhibitors C75 and G28UCM were applied either with or following NMDA. Retinal sections were labeled for EdU (red) and Sox2 (green; e), TUNEL (h), or DRAQ5 (magenta), EdU (red) and CD45 (green; **k**,**l**). Arrows indicate nuclei of MG, small double-arrows nuceli of NIRG cells and hollow arrow-heads indicate nuclei of microglia. The histograms **f**, **g**, **i**, **j** and m-p represents the mean (± SD) and each dot represents one biological replicate retina. The calibration bar in **c** represents 50 µm. Abbreviations: ONL – outer nuclear layer, INL – inner nuclear layer, IPL – inner plexiform layer, GCL – ganglion cell layer. Significant changes (*p<0.01, **p<0.001, ***p<0.0001) in expression were determined by using a Wilcox rank sum with Bonferroni correction.

Treatment of NMDA-damaged retinas with FASN inhibitors, G28UCM and C75, resulted in significant reductions in numbers of proliferating MGPCs (Fig. 11e-g). These large decreases in numbers of proliferating MGPCs occurred despite significant decreases in cell death (Fig. 11h-j) and increases in proliferation of microglia (Fig. 11k-p) in NMDA-damaged retinas treated with FASN inhibitors. Increases in numbers of proliferating microglia occurred without increasing total numbers of CD45+ cells (Fig. 11k,p), suggesting that there was less recruitment of peripheral monocytes coincident with increased proliferation of resident microglia to result in no net change in total numbers of CD45+ cells in the retina. NIRG cells were not significantly affected by FASN inhibitors (not shown).

We next sought to investigate whether cell signaling in damaged retinas was influenced by FASN-inhibitor. We observed a significant decrease in levels of pS6 in the cytoplasm of MG treated with FASN-inhibitor in damaged retinas (Fig. 12a,b). By comparison, levels of pStat3 and pSmad1/5/8 were undetectable in the nuclei of MG treated with FASN-inhibitor in damaged retinas (n≥5) (Fig.12a). Signaling though mTor (pS6), Jak/Stat- and BMP/SMAD-signaling are known to be required for the formation of proliferating MGPCs (Todd et al., 2016a; Todd et al., 2017; Zelinka et al., 2016). Inhibition of FASN in NMDA-damaged retinas had no effect upon numbers of TUNEL-positive dying cells (Fig. 12a,d).

**Figure 12.**
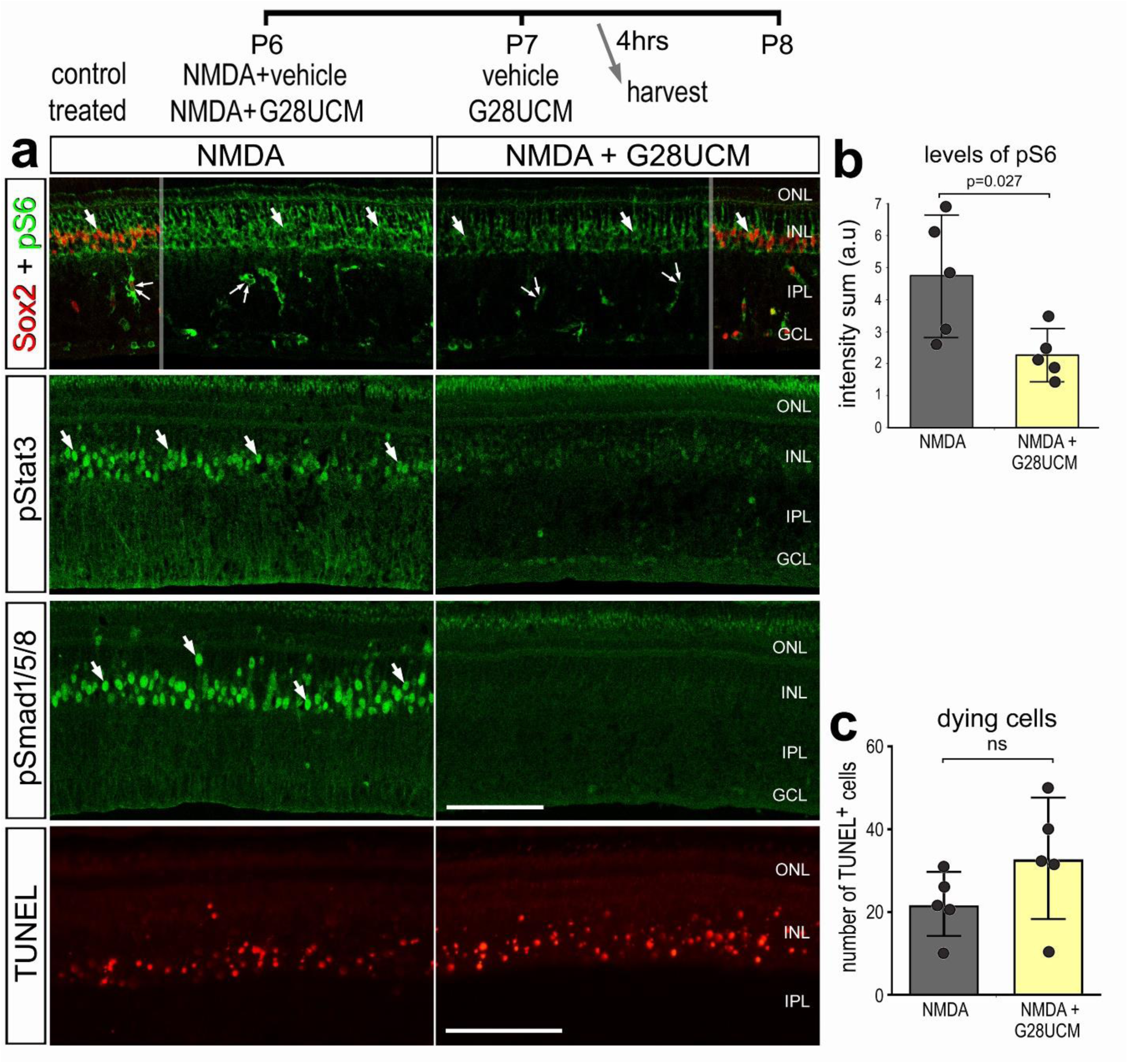
Inhibition of FASN suppresses cell signaling in MG. Retinas were obtained from eye injected with NMDA ± G28UCM (FASN inhibitor) at P7, vehicle ± G28UCM and harvested 4 hours after the last injection. Retinal sections were labeled for Sox2 (red) and pS6 (green), pStat3 (green), pSmad1/5/8 (green), and fragmented DNA in the nuclei of dying cells (**a**). Arrows indicate the nuclei of MG. The calibration bar in **a** represents 50 µm. Abbreviations: ONL – outer nuclear layer, INL – inner nuclear layer, IPL – inner plexiform layer, GCL – ganglion cell layer. The histograms represent the mean (± SD) fluorescence intensity sum (a.u.; **b**) or mean number of TUNEL+ cells (**c**) and each dot represents one biological replicate retina. Significance (p-values) of difference was determined by using a two-tailed paired Student’s t-test.

## Discussion

In this study we investigate the function of FASN and FABPs in glial cells in the chick retina. We observed that *FABP7* is highly expressed by eRPCs and maturing MG during embryonic development. *FABP7, FABP5* and *PMP2* are up-regulated during the activation of MG after damage or treatment with FGF2 and insulin. This pattern of FABP expression was also observed for NIRG cells and microglia in damaged retinas. Inhibition of FABPs or FASN influenced the proliferation of different types of cells including NIRG cells, microglia, and MGPCs. Inhibition of FABPs in undamaged retinas induced reactivity in MG, but also both decreased levels of genes associated with resting mature MG and neurogenesis and increased genes associated with gliogenesis and inflammation. Inhibition of FABPs and FASN in damaged retinas selectively suppressed cell signaling pathways in MG that are known to promote the formation of MGPCs. These findings indicate the importance of FASN and FABPs in mediating the transition into a proliferating MGPCs.

### FABPs in retinal development

Different FABP isoforms are known to be expressed by different cell types in maturing mammalian brain (Owada, 2008). Similarly, FABP isoforms have been identified in the chick retina (Sellner, 1993). FABP7 is often used as a biomarker for radial glia (brain lipid binding protein, Blbp) in the developing mouse brain and is presumed to facilitate cortical development (Anthony et al., 2005; Feng et al., 1994). FABPs are expressed in different types of tumors, particularly in cancer metastasis (Ohmachi et al., 2006; Senga et al., 2018). We found that *FABP7* was upregulated in developing and maturing MG in embryonic chick retina. In addition, *FABP7* was detected in developing amacrine and bipolar interneurons, but was downregulated in mature neurons. By comparison, *FABP5* was expressed at high levels in mature interneurons. These cell-type specific patterns of expression for FABPs may indicate isoform-specific rolls despite overlap in ligand-binding among FABPs.

The chick eye is known to have proliferating retinal progenitor cells at the ciliary marginal zone of the retina (Fischer and Reh, 2000; Fischer et al., 2014a). Inhibition of FABPs did not influence the proliferation of CMZ progenitors. These progenitors expressed relatively low levels of PMP2, but expression of FABP5 and FABP7 in these cells remains unknown. Treatment with insulin and FGF2 upregulated PMP2 in MG and CMZ progenitors, yet FABP inhibitor did not influence the proliferation of the CMZ progenitors. It is possible that FABP inhibitor had no effect because the drug failed to diffuse through the vitreous to act at the CMZ. Alternatively, FABPs do influence the proliferation of CMZ progenitors. Although many cell signaling pathways can influence both CMZ progenitors and MGPCs, there are instances where CMZ progenitors and MGPCs are differentially influenced by cell signaling pathways. For example, insulin and IGF1 stimulate the proliferation of CMZ progenitors (Fischer and Reh, 2000; Fischer and Reh, 2002), whereas the insulin and IGF1 must be combined with FGF2 to stimulate the proliferation of MGPCs (Fischer et al., 2002; Ritchey et al., 2012). Similarly, HG-EGF stimulates the proliferation of MGPCs, but has no effect upon CMZ progenitors even when combined with IGF1 (Todd et al., 2015). By comparison, glucagon suppresses the proliferation of CMZ progenitors (Fischer et al., 2005), whereas glucagon has no effect upon the proliferation of MGPCs (unpublished observations). By contrast, the proliferation of CMZ progenitors and MGPCs can be stimulated by Sonic Hedgehog signaling (Moshiri et al., 2005; Todd and Fischer, 2015), stimulated by retinoic acid (Todd et al., 2018) and by inhibition of Smad3 (Todd et al., 2017).

### NIRG and Oligodendrocyte proliferate after damage

During embryonic development glial precursor cells migrate into the retina from the optic nerve (Rompani and Cepko, 2010). These precursor cells undergo a cell division to generate oligodendrocytes and diacytes (Rompani and Cepko, 2010), also known as non-astrocytic inner retinal glia (NIRG) cell that reside predominantly in the IPL (Fischer et al., 2010). This unique type of glial cell has been identified in the retinas of birds and some types of reptiles (Todd et al., 2016b). The functions of the NIRG cells is unknown, but these cells are known to proliferate in response to IGF1 (Fischer et al., 2010) and their survival is tethered to retinal microglia (Zelinka et al., 2012b). We find that NIRG cells express PMP2, but only in peripheral regions of the retina or at later times (>7 days) after NMDA-induced damage. By comparison, inhibition of FASN had no effect upon the proliferation of NIRG cells.

Consistent with previous reports (Kohsaka et al., 1980; Kohsaka et al., 1983), the chicken retina contains axons that are thinly myelinated by oligodendrocytes that express Sox10, Olig2 and PMP2. Surprisingly, we observed that these oligodendrocytes express HuC/D, a common neuronal marker of amacrine cells and ganglion cells. Thus, unambiguous identification of neurons in the GCL requires markers in addition to HuC/D. There were rare PMP2^+^ cell in the inner INL of peripheral regions of retina that did not express neuronal markers, but expressed a set of markers associated with NIRG cells. One week after damage there was an increase in the number of EdU-labeled oligodendrocytes, which suggests *de novo* myelination of axons in the NFL from newly generated oligodendrocytes. Further studies are required to determine whether the newly generated oligodendrocytes directly results in additional axon myelination. Notably, NMDA-damage is not expected to result in demyelination, which raises questions about the signals that promote the proliferation of oligodendrocytes. Without genetic models or viruses with appropriate trophism to lineage-trace the origin of newly generated oligodendrocytes, the origin of oligodendrocyte precursor cells remains uncertain.

### The formation of proliferating MGPCs requires FABPs and FASN

scRNA-seq data indicate that *FABP7* is expressed by resting MG in the postnatal chick retina. When the retina is damaged or treated with FGF2 and insulin the MG robustly upregulate *FABP5, FABP7* and *PMP2*. When FABPs are inhibited in damaged retinas, significantly fewer proliferating MGPCs are generated (Hoang et al., 2020). Similarly, MGPC proliferation was suppressed when inhibitor was applied before damage when only FABP7 is highly expressed in MG. We observed diminished cell signaling through pSMAD1/5/8, pStat3, and mTor (pS6) in MG treated with FABP inhibitor. These findings are consistent with treatment-specific Ligand-Receptor interactions in glia treated by FABP inhibitor revealed by SingleCellSignalR analyses. Similarly, levels of pSMAD1/5/8 and pStat3 fell below detection in MG treated with FASN-inhibitor. These cell signaling pathways have been shown to promote the formation of proliferating MGPCs in the chick retina (Todd et al., 2016a; Todd et al., 2017). Further, we find loss of Ligand-Receptor interactions involving FGF, Midkine and Notch1 in retinal glia treated with FABP inhibitor. (Anthony et al., 2005). These pathways are necessary the formation of proliferating MGPCs (Fischer et al., 2009a; Fischer et al., 2009b; Ghai et al., 2010; Hayes et al., 2007). Consistent with our observations, Notch-signaling is known to be downstream of FABPs (Anthony et al., 2005).

FABP isoforms serve many different functions including cellular metabolism and cellular trafficking of lipid metabolites (Storch and Corsico, 2008). In addition to facilitating lipid metabolism, FABPs can also facilitate nuclear transport of hydrophobic ligands for cell signaling such as PPAR (Tripathi et al., 2017), retinoic acid (Dawson and Xia, 2012), and endocannabinoids (Haj-Dahmane et al., 2018). Given the well-established involvement of FABPs in lipid metabolism, we found associations with the expression of FASN, which is important for producing long chain fatty acids (Kuhajda, 2006). When we antagonize FASN with different small molecule inhibitors there are significant decreases in numbers of proliferating MGPCs. These findings provide further evidence that lipid metabolism is required for the transition of resting MG to activated states, and subsequent proliferation as progenitor-like cells. scRNA-seq data indicate that MG become reactive with FABP inhibition in the absence of neuronal damage; there was significant upregulation of genes associated with gliogenesis and reactivity. This broad shift in the gene-expression modules to induce reactive phenotypes was supported by evidence for Ligand-Receptor interactions associated with reactivity including signals such as IL1β, TGFβ and HB-EGF.

Although inhibition of FABPs in undamaged retinas stimulated MG to adopt a reactive phenotype and acquire a transcriptomic profile characteristic of activation and de-differentiation, FABP-inhibition prior to NMDA did not prime MG to become MGPCs. This likely resulted from FABP-inhibition up-regulating gene modulates associated with gliogenesis and down-regulation of gene modulates associated with neurogenesis and proliferation. FABP-inhibition in undamaged retinas should have inhibited FABP7 in resting MG and PMP2 in oligodendocytes, and this inhibition may have resulted in the activated transcriptomic profiles seen in MG and microglia. However, treatment with FABP inhibitor resulted in an upregulation of *FABP5* and *PMP2* in MG and *FABP5, FABP7* and *PMP2* in microglia. Thus, it is possible that the suppressed formation of MGPCs following FABP inhibition resulted from upregulation and inhibition of FABPs in MG and microglia.

### FABP-inhibition suppresses proliferation and induces reactivity in microglia

The development of FABP inhibitors was motivated by the presence of FABP4 in obese patients suffering from atherosclerosis, where macrophages contribute the narrowing of arterial vessels (Furuhashi et al., 2007; Makowski et al., 2001). Peripheral macrophages express FABP4 and inhibition may slow the progression of vessel narrowing (Furuhashi et al., 2007; Makowski et al., 2001). After retinal damage in chick, microglia normally proliferate and acquire a reactive phenotype (Zelinka et al., 2012b). The presence of reactive microglia is required for the formation of MGPCs (Fischer et al., 2014b). Further, signals from reactive microglia mediate inflammatory signaling in MG through pathways such as NFkB (Palazzo et al., 2020). Given that microglia rapidly respond to retinal damage with upregulation of pro-inflammatory signals (Todd et al., 2019), it is possible that reduced numbers of microglia in damaged retinas treated with FABP inhibitor also influenced the formation of proliferating MGPCs. In damaged retinas treated with FABP inhibitor, microglia appeared highly reactive, but the numbers of these cells were significantly reduced. Further, we did not detect significant numbers of microglial genes that were differentially expressed in damaged retinas treated with FABP inhibitor. Thus, it seems most likely that microglia did not influence the formation of MGPCs in inhibitor-treated retinas.

## Conclusions

FASN and FABPs are novel targets of investigation with respect to retinal glia and reprogramming of MG into MGPCs. We found that FABPs are highly expressed by MG during reprogramming into proliferating MGPCs. Inhibition of FABPs results in the upregulation of genes associated with gliogenesis and inflammation while concurrently reducing the expression of genes associated with proliferation and neurogenesis. The anti-proliferative effects of FABP inhibition were not specific to MG, as microglia also showed reduced proliferation in inhibitor-treated retinas. By contrast, the proliferation of CMZ was unaffected by FABP inhibitor. Our findings suggest that FABPs mediate glial reactivity and de-differentiation through lipid-associated cell signaling, while proliferation requires lipid metabolism. Consistent with this hypothesis, inhibition of FASN potently inhibited the formation of proliferating MGPCs, while decreasing cell death and increasing microglial proliferation. Microglia express FABPs, and FABP inhibition alters cytokine production and reactivity which is expected to impact signaling with MG. Collectively, our data suggest the activity of FASN and FABPs MG to become activated prior to forming proliferating MGPCs in the chick retina.

## Author contributions

WAC – experimental design, execution of experiments, collection of data, data analysis, construction of figures and writing the manuscript. AT, EH and MH – execution of experiments and collection of data. HE - experimental design, collection of data, data analysis and writing the manuscript. TH and SB – preparation of scRNA-seq libraries. AJF – experimental design, data analysis, construction of figures and writing the manuscript.

## Data availability

RNA-Seq data for gene-cell matrices are deposited at GitHub https://github.com/jiewwwang/Single-cell-retinal-regeneration https://github.com/fischerlab3140/scRNAseq_libraries

Some of the scRNA-seq data can be queried at https://proteinpaint.stjude.org/F/2019.retina.scRNA.html.

## Acknowledgements

This work was supported by RO1 EY032141- 01 (AJF) and UO1 EY027267-04 (AJF, SB).

---

**Supplemental Figure 1.**
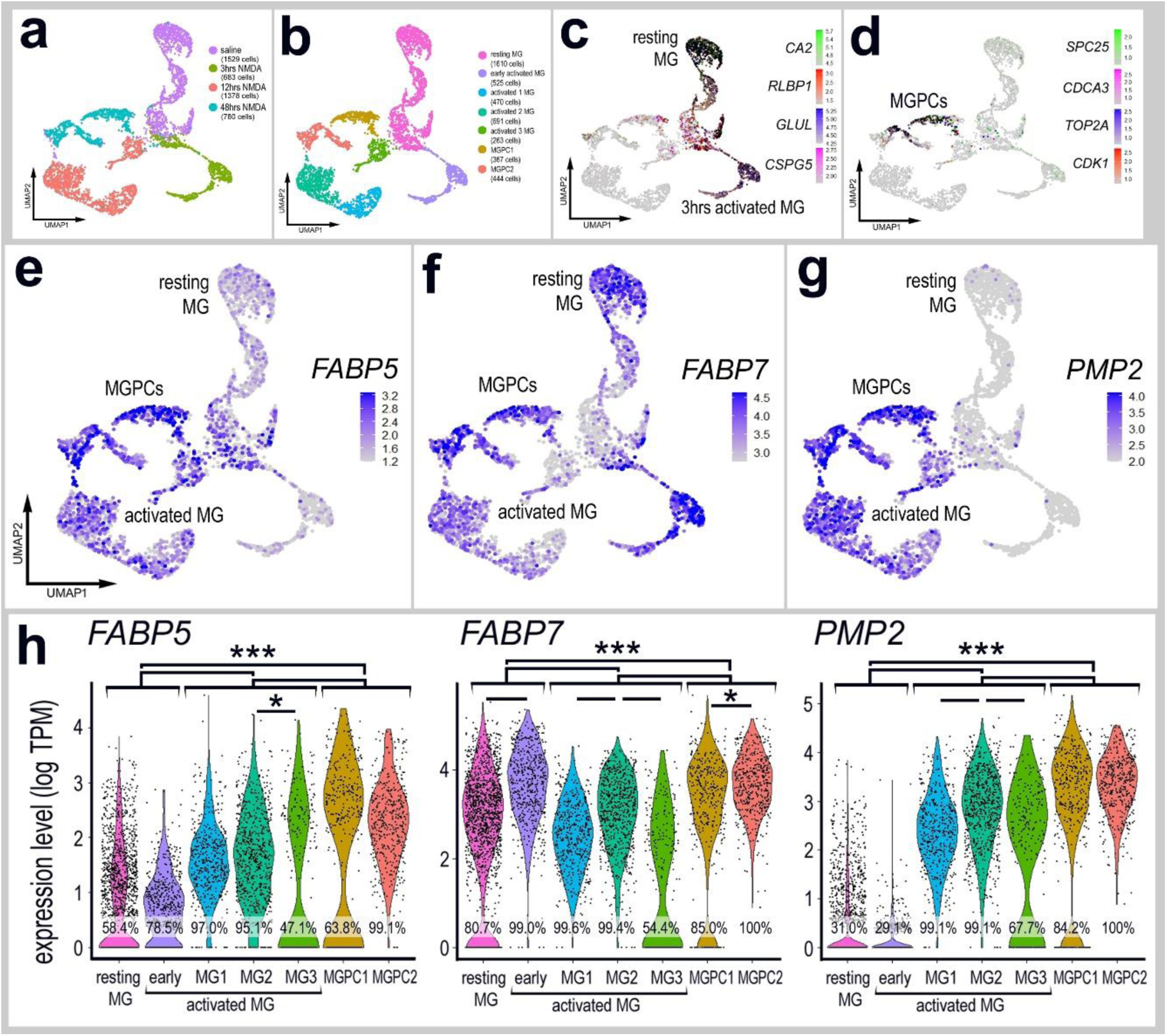
scRNA-seq was used to interrogate expression of FABPs in MG at timepoints soon after damage. Retinas were harvested at 3, 12 or 48 hrs after treatment with NMDA. UMAP plots illustrate the distribution of different libraries (saline, 4hrs, 12hrs and 48hrs after NMDA) (**a**). UMAP ordering of cells revealed 7 different clusters of cells (**b**). Resting MG were identified based on elevated levels of expression for *CA2, RLBP1, GLUL* and *CSPG5*, whereas activated MG down-regulated these markers (c). MGPCs were identified based on expression of proliferation markers including *SPC25, CDCA3, TOP2A* and *CDK1* (**d**). UMAP heatmap plots illustrate patterns of expression of *FABP5* (**e**), *FABP7* (**f**) and *PMP2* (**g**). Expression levels (log TPM) and the percentage of expressing MG are illustrated in violin plots (**h**). Significance (*p<0.01, ***p<0.0001) of difference in expression was determined by using a Wilcox rank sum with Bonferroni correction.

**Supplemental Figure 2.**
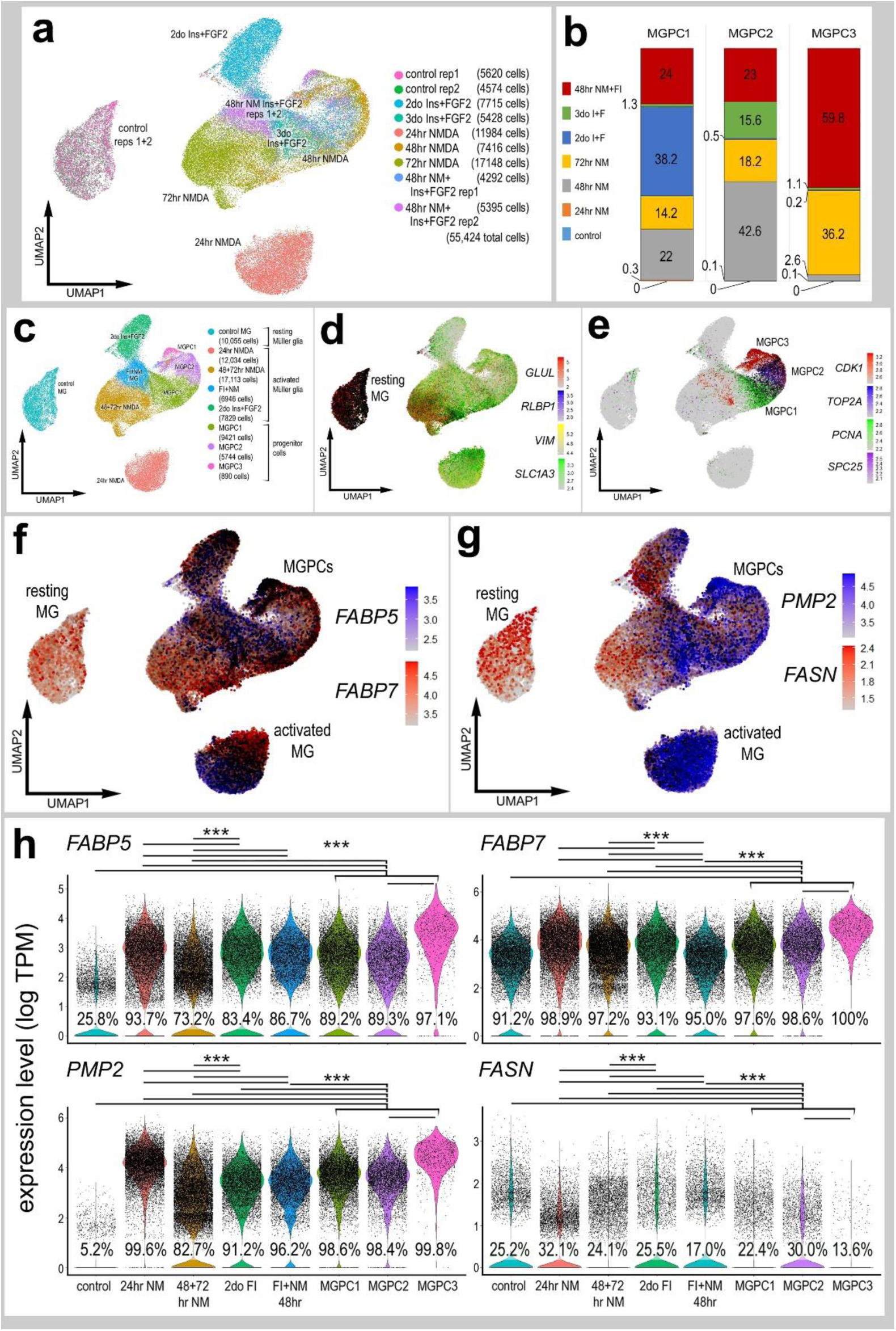
Comparison of *FABP* and *FASN* expression levels in MG and MGPCs from different treatment conditions. scRNA-seq was used to identify patterns of expression of *FABPs* and *FASN* in MG and MGPCs at different time points after NMDA damage and/or FGF + insulin growth factor treatment. Nine different libraries were aggregated for a total of more than 55,000 MG and MGPCs (**a-c**). MGPCs were identified based on high-levels of expression of *CDK1, TOP2A, PCNA* and *SPC25* (**e**), and were a mix of cells from 48hrs NMDA+FGF2+insulin, 48hrs NMDA, 72hrs NMDA and 3 doses of insulin+FGF2 (**b**). Resting MG were identified based on expression of *GLUL, RLBP1, SLC1A3* and *VIM* (**c,d**). Each dot represents one cell and black dots indicate cells with 2 or more genes expressed. The expression of *FABP5, FABP7, PMP2* and *FASN* is illustrated UMAP and violin plots with population percentages and statistical comparisons (**f-h**). Significance of difference (***p<10-20) in expression levels (log TPM) were determined by using a Wilcox rank sum with Bonferoni correction. Abbreviations: ONL –outer nuclear layer, INL –inner nuclear layer, IPL –inner plexiform layer, GCL –ganglion cell layer.

**Supplemental Figure 3.**
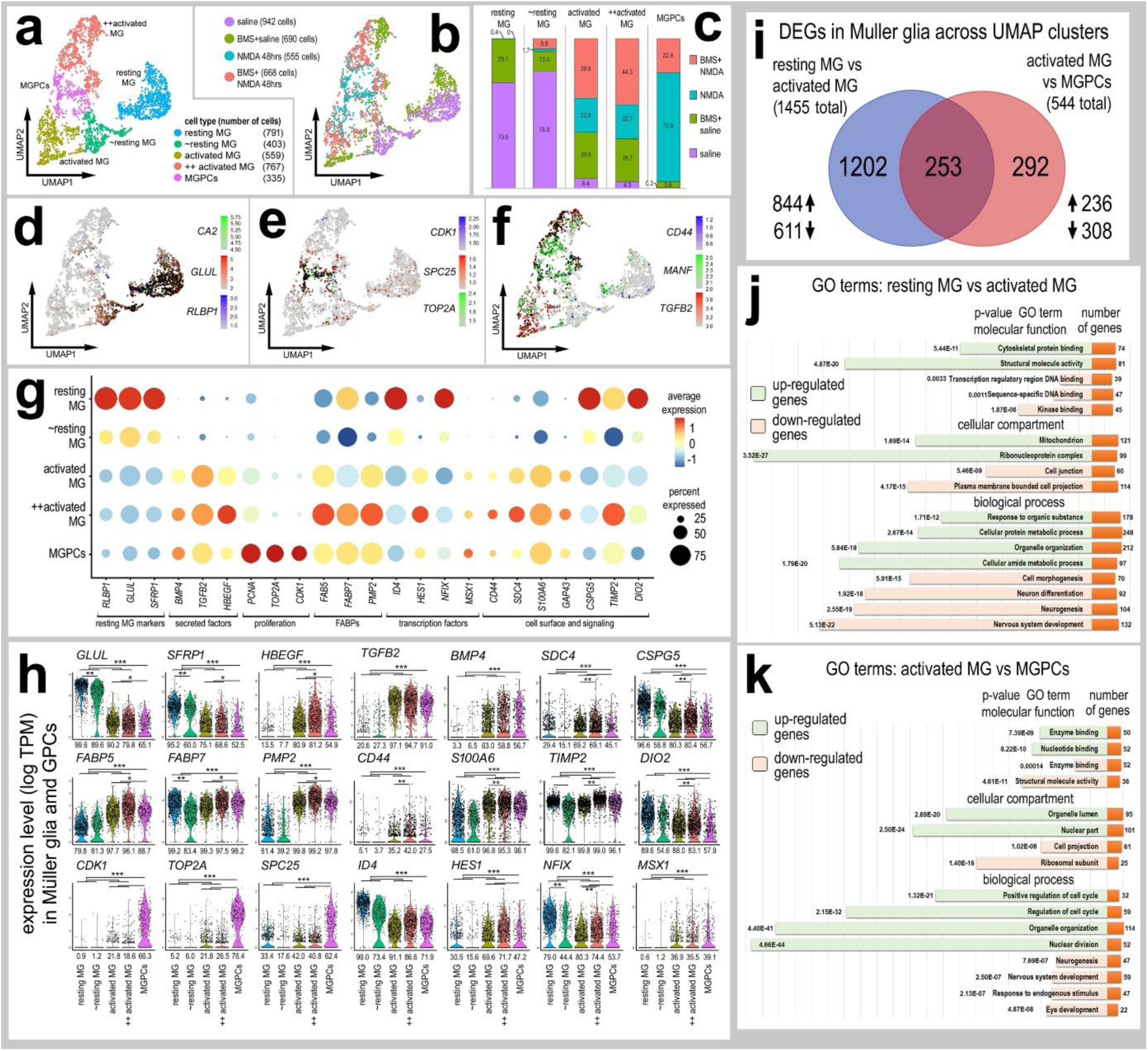
scRNA seq libraries were generated to analyze changes in MG gene expression. MG were identified based on expression of genes associated with resting glia, activated glia, and proliferating MGPCs. UMAP ordering of MG revealed 2 clusters of resting MG, 2 clusters of activated MG, and one cluster of MGPCs (**a**). Resting MG clusters were predominantly occupied by MG from saline-treated retinas, activated MG clusters were occupied by cells treated with saline-BMS, NMDA alone and NMDA-BMS, and the MGPC cluster was predominantly occupied by cells from NMDA-treated retinas (**b**,**c**). Resting MG were identified based on expression of markers such as *CA2*, *GLUL* and *RLBP1* (**d**). MGPCs were identified based on expression of proliferation markers such as *CDK1*, *SPC25* and *TOP2A* (**e**). Activated MG were identified based on expression of markers such as *CD44, MANF* and *TGFB2* (**f**). Differentially expressed genes (DEGs) were identified for MG from retinas treated with saline vs BMS-saline, saline vs NMDA, and NMDA vs BMS-NMDA and plotted in a Venn diagram (**i**). The Dot plot plots indicating the percentage of expressing MG (size) and expression levels (heatmap) for genes related to resting glia, secreted factors, glial transcription factors, inflammation, glial reactivity and proliferation (**g, h**). All genes displayed in the Dot plot have significantly different (p<0.0001) expression levels in MG from retinas treated with saline vs saline-BMS (**g**). Gene Ontology (GO) terms for the enriched genes in the BMS treated and BMS+NMDA treated were compiled (ShinyGO) and grouped by biological process, cellular component and molecular function. GO enrichment analysis was performed for up-regulated DEGs (light green) and down-regulated DEGs (light orange). The significance of the GO category and the number of enriched genes grouped into each function (orange) are displayed.

## Notes

### Competing Interest Statement

The authors have declared no competing interest.

